# Structural basis of closed groove scrambling by a TMEM16 protein

**DOI:** 10.1101/2023.08.11.553029

**Authors:** Zhang Feng, Omar E. Alvarenga, Alessio Accardi

**Author notes:** correspondence to: Alessio Accardi.

## Abstract

Activation of Ca^2+^-dependent TMEM16 scramblases induces the externalization of phosphatidylserine, a key molecule in multiple signaling processes. Current models suggest that the TMEM16s scramble lipids by deforming the membrane near a hydrophilic groove, and that Ca^2+^ dependence arises from the different association of lipids with an open or closed groove. However, the molecular rearrangements involved in groove opening and of how lipids reorganize outside the closed groove remain unknown. Using cryogenic electron microscopy, we directly visualize how lipids associate at the closed groove of Ca^2+^-bound nhTMEM16 in nanodiscs. Functional experiments pinpoint the lipid-protein interaction sites critical for closed groove scrambling. Structural and functional analyses suggest groove opening entails the sequential appearance of two π-helical turns in the groove-lining TM6 helix and identify critical rearrangements. Finally, we show that the choice of scaffold protein and lipids affects the conformations of nhTMEM16 and their distribution, highlighting a key role of these factors in cryoEM structure determination.

## Introduction

Activation of phospholipid scramblases disrupts the plasma membrane asymmetry, and results in the exposure of the negatively charged lipids phosphatidylserine (PS), an important signaling molecule. Thus, lipid scrambling plays a key role in a variety of physiological processes, ranging from recognition of apoptotic cells by macrophages and blood coagulation, to viral entry, membrane fusion and repair ^1–3^. To date, three families of integral membrane proteins have been identified as lipid scramblases, the Ca^2+^-activated TMEM16s ^4–7^, the caspase-dependent XKRs ^8^, and the ATG9 proteins involved in autophagosome formation ^9–11^. Additionally, several other proteins and single-helix membrane spanning peptides have been reported to mediate lipid scrambling ^12–16^. These scramblases share broadly conserved functional characteristics, such as mediating rapid and poorly selective lipid transport ^1–3,6,7^, while displaying a remarkable structural diversity, as each family adopts a different fold ^7,9–11,17–22^. Thus, it is not known whether these diverse scramblases function according to a conserved mechanism.

The currently accepted paradigm for lipid scrambling, the so-called credit-card mechanism, is that scramblases provide a hydrophilic pathway that shields the polar lipid headgroups as they traverse the bilayer, while the acyl chains remain embedded in the hydrocarbon core of the membrane ^7,23^. In several dimeric TMEM16 scramblases, Ca^2+^ binding favors opening of a transmembrane hydrophilic groove which has been proposed to serve as the translocation pathway for lipid headgroups ^7,24–28^. The lipid groove is formed by TM3-7 helices of each TMEM16 protomer, and its interior is lined by hydrophilic residues ^7^ (Fig. 1a-c and f-h). In the apo conformation, the TM4 and TM6 form an extensive interface that seals the groove interior from the hydrocarbon core of the membrane (Fig. 1a, f), resulting in a closed groove. The Ca^2+^ binding sites are located between TM6-8 and their formation is enabled by a rearrangement of the intracellular portion of TM6 (Fig. 1b, g). Groove opening follows Ca^2+^ binding and entails a movement of the TM3 and TM4 helices away from TM6 ^17,18,21^ (Fig. 1c, h). However, Ca^2+^ binding does not promote groove opening in all TMEM16 scramblases as the mammalian TMEM16F exclusively adopts a Ca^2+^-bound closed groove conformation, even in conditions of maximal activity ^19,22,29^, leading to the proposal that the closed groove is also a scrambling competent state ^19,30,31^. Gain-of-function mTMEM16F mutants adopt conformations where the groove remains closed, but with a rearranged TM4-6 interface ^29^, raising the possibility that different closed-groove conformations might be scrambling competent. Thus, how Ca^2+^ binding to the TM6-lined sites induces the rearrangement of TM3 and TM4 to induce groove opening in some but not all TMEM16s is not known.

**Figure 1.**
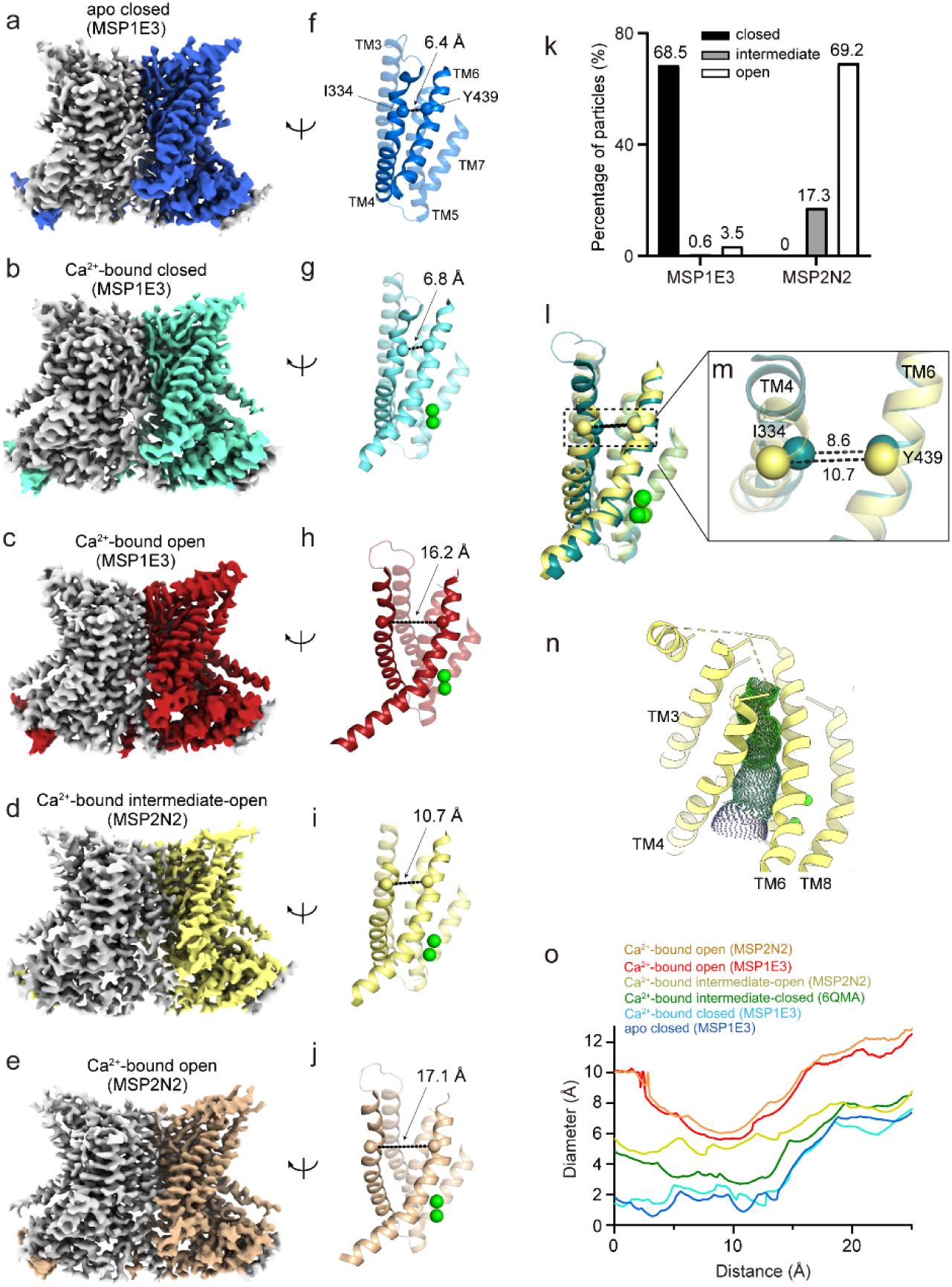
Cryo-EM structures of nhTMEM16 in the MSP1E3 and MSP2N2 nanodiscs. **a**-**b**, Cryo-EM maps of nhTMEM16 in nanodiscs formed from a 7:3 mix of DOPC:DOPG lipids and MSP1E3 (**a**-**c**) or MSP2N2 (**d**-**e**) scaffold proteins. (**a**) Ca^2+^-free, closed groove, (**b**) Ca^2+^-bound, closed groove, (**c**) Ca^2+^-bound, open groove, (**d**) Ca^2+^-bound, intermediate-open groove, and (**e**) Ca^2+^-bound, open groove. In all dimers, one protomer is colored in gray and the other in blue (**a**), cyan (**b**), red (**c**), yellow (**d**), and sand (**e**). **f-j**, The groove in the different conformations is viewed from the plane of the membrane. Transmembrane helices are shown in cartoon representations and labelled. States are colored as in (**a**-**e**). Colored spheres correspond to the position of the Cα atoms of I334 on TM4 and Y439 on TM6 and their distance is indicated. Ca^2+^ ions are shown as green spheres. **k**, The percentage of particles with a closed (black bar), intermediate (gray bar), or open (white bar) groove conformation in the datasets of Ca^2+^-bound nhTMEM16 in MSP1E3 or MSP2N2 nanodiscs. Here, intermediate does not distinguish between the intermediate-open and -closed conformations. **l-m**, Alignment of the groove helices in the intermediate-open (yellow) and previously reported intermediate-closed (PBD: 6QMA, dark green) states (**l)**, the distance between the Cα atoms of I334 on TM4 and Y439 on TM6 increases by ∼2 Å in the intermediate-open **(m)**. **n**, The accessibility of the permeation pathway of nhTMEM16 in the intermediate conformation is visualized using the program Hole ^39^. **o**, The inner diameter of the permeation pathway of nhTMEM16 in MSP1E3 or MSP2N2 nanodiscs and in the published the diameter of a putative ion conduction pathway measured in different states with Hole including the Ca^2+^-bound open state in MSP2N2 (orange), Ca^2+^-bound open state in MSP1E3 (red), Ca^2+^-bound intermediate-open state in MSP2N2 (yellow), Ca^2+^-bound closed state in MSP1E3 (cyan), the previously reported intermediate-closed state in MSP2N2 (dark green) and Ca^2+^-free closed in MSP1E3 (blue).

Cryogenic electron microscopy (cryoEM) imaging of TMEM16 scramblases in nanodiscs showed these proteins induce a pronounced thinning of the membrane near the groove region, a distortion that is more pronounced when the groove opens ^17,21,30^. However, resolution of these structures was insufficient to visualize how individual lipids interact with the groove in the open and closed conformations. Recently, high-resolution structures of the Ca^2+^-bound fungal afTMEM16 with an open groove revealed that lipids adopt distorted poses with their headgroups positioned outside the pathway, providing the bases for membrane thinning near the open groove ^30^. Further, functional experiments suggested that specific interactions between groove-lining residues and lipid headgroups are not required for open groove scrambling ^30^. Together, these observations led to the proposal that scrambling at an open groove is enabled by membrane thinning ^30^, rather than occurring via a credit card mechanism. Because membrane thinning is also observed in conditions when the groove is closed, it was proposed that afTMEM16 ^30^ and TMEM16F ^19,30,31^ could scramble lipids outside a fully closed groove, or one with a rearranged TM4-TM6 interface ^29^. This proposal could also account for the basal activity of many TMEM16 scramblases in the absence of Ca^2+^ when the groove is closed ^5,7,18^. However, direct structural information on how lipids arrange near a closed groove is lacking, and it is not known whether or how their arrangement plays a role in closed groove scrambling.

One important factor to consider when interpreting cryoEM imaging experiments is the potential role of the environment in determining the conformation and state distribution of proteins. For example, the Ca^2+^-bound human TMEM16K scramblase crystalizes with an open groove but has a closed groove in single particle cryoEM imaging conditions ^18^, and the fungal nhTMEM16 adopts conformations with an open, closed, and intermediate-closed groove with roughly equal probability when reconstituted in MSP2N2 nanodiscs but is only in an open groove state when imaged in detergent micelles with cryoEM and x-ray crystallography ^7,21^. In contrast, other homologues, such as murine TMEM16F and fungal afTMEM16 are less sensitive to the environment, as they exclusively adopt a single conformation in detergent and in nanodisc environments ^17,19,22,30^. Thus, the extent to which choices of detergent, lipid and/or nanodisc scaffold protein influence the conformation of TMEM16 scramblases is poorly understood.

Here, we used cryoEM to image the fungal nhTMEM16 scramblase in lipid nanodiscs formed from different scaffold proteins and found this to affect both the conformations adopted by nhTMEM16 and their distribution. In the smaller MSP1E3 nanodiscs the scramblase preferentially adopts a closed conformation, whereas in the larger MSP2N2 nanodiscs nhTMEM16 predominantly adopts open and intermediate-open conformations. The 2.64 Å resolution structure of Ca^2+^-bound closed nhTMEM16 reveals how lipids arrange near the groove, thus showing how the scramblase thins the membrane outside a closed groove. Mutating residues that coordinate outer leaflet lipids near the closed groove impairs scrambling only in the absence of Ca^2+^, suggesting these interactions play a specific role in closed-groove scrambling. Structural analysis of the transition of nhTMEM16 from closed to open state suggests that the E313-R432 salt bridge could be involved in groove opening. Disruption of this interaction by the R432A mutation trapped nhTMEM16 in a closed-groove conformation, suggesting the salt bridge is critical for groove opening. In sum, our work provides insights into how lipids are scrambled outside a closed groove, how the groove opens, and how these processes are regulated by the lipid environment.

## Results

### The conformational landscape of nhTMEM16 depends on the environment

To understand how the environment affects the conformational landscape of TMEM16 scramblases, we focused on the fungal nhTMEM16 as this homologue can adopt multiple conformations in different reconstitution systems ^7,21^. We imaged the scramblase in the absence and presence of Ca^2+^ and in nanodiscs formed from a 7:3 mixture of 1,2-Dioleoyl-sn-glycero-3-phosphocholine (DOPC) and 1,2-Dioleoyl-sn-glycero-3-phosphatidylglycerol (DOPG) lipids. We used the MSP1E3 and/or MSP2N2 scaffold proteins that differ in their average diameter by ∼3 nm ^32,33^. In 0 Ca^2+^ and in MSP1E3 nanodiscs, we obtained a single class of particles, which yielded a map with average resolution of 2.93 Å corresponding to a conformation with a closed hydrophilic groove and no density in the Ca^2+^ binding sites (Fig. 1a, f and Extended Data Fig. 1). While this is similar to what was reported for nhTMEM16 in nanodiscs formed with the larger MSP2N2 and POPC:POPG lipids, there are differences in the Ca^2+^ sites and position of the cytosolic domains (see below). In contrast, the conformational distribution of nhTMEM16 changes drastically when reconstituted in 0.5 mM Ca^2+^ in MSP1E3 or MSP2N2 nanodiscs. We found that in MSP2N2 nanodiscs the majority of the particles (∼69.2%) adopt an open groove conformation, ∼17.3% are in an intermediate-open state (see below), and no class with a closed groove could be detected (Fig. 1d-e, i-j and Extended Data Fig. 2). The remaining ∼13.5% particles have poorly defined TM4 helices and could not be assigned (Extended Data Fig. 2). This is different from the findings in MSP2N2 POPC/POPG nanodiscs ^21^, where the closed groove conformation was also significantly populated. In the smaller MSP1E3 nanodiscs nhTMEM16 predominantly adopts a closed groove conformation (∼68.5% of particles) (Fig. 1b, g and Extended Data Fig. 3) and only a small fraction is in an open groove conformation (∼3.5% of particles) (Fig. 1c, h and Extended Data Fig. 3). In the remaining ∼28% of particles the density around the groove region was not well resolved, suggestive of conformational heterogeneity. Classification of the symmetry-expanded and signal-subtracted protomers from this subset shows that ∼16% adopt a conformation with a closed groove, ∼15% are in an intermediate conformation and ∼11% have an open groove (Extended Data Fig. 3 and 4). In the remaining ∼58% protomers the density for TM4 is of insufficient quality for model building but is consistent with the groove being in closed or intermediate conformations (Extended Data Fig. 4). Mapping these protomers to the original dimeric particles allowed for a reconstruction of a nhTMEM16 dimer with one open and one closed groove (Extended Data Fig. 3), consistent with the idea that the protomers can gate autonomously ^34,35^. In sum, these results show that the open groove conformation of nhTMEM16 is favored >20-fold by reconstitution in the larger MSP2N2 nanodiscs (Fig. 1k).

Next, we analyzed whether the choice of nanodisc scaffold protein and lipid composition also affect the structures of the observed nhTMEM16 conformations. Unexpectedly, we identified several structural differences. The Ca^2+^-free map of nhTMEM16 determined in MSP1E3 DOPC/DOPG nanodiscs (PBD: 6QM4, EMD: 4589) ^21^ differs from the previously determined one in MSP2N2 POPC/POPG nanodiscs as the Ca^2+^ binding sites are empty (Extended Data Fig. 5a, d), TM6 is bent, and the cytosolic domains move away from each other by ∼4 Å (Extended Data Fig. 5a). This conformation, with a bent TM6, empty Ca^2+^ sites, and separated cytosolic domains, resembles that of Ca^2+^-free afTMEM16 ^17,30^. Indeed, the conformation of nhTMEM16 in 6QM4 is structurally more similar to its Ca^2+^-bound closed state (Cα r.m.s.d of 0.67 Å, Extended Data Fig. 5b) than to our apo conformation (Cα r.m.s.d. 1.76 Å, Extended Data Fig. 5c). These rearrangements are likely due to the presence of an unassigned density in the Ca^2+^ binding sites in the EMD-4589 map (Extended Data Fig. 5e). The identity of the ligand causing the observed rearrangements is unknown. This suggests that ligand binding to one of two sites is sufficient to induce a rearrangement in the cytosolic domains and TM6.

The Ca^2+^-bound open conformation determined in MSP1E3 DOPC/DOPG nanodisc is nearly identical to the previously determined open structure of nhTMEM16 determined in the larger MSP2N2 POPC/POPG discs (PDB: 6QM9) ^21^ (Cα r.m.s.d ∼0.58 Å, Extended Data Fig. 5h). The open conformation in MSP2N2 DOPC/DOPG nanodiscs has a slightly more opened groove than that in POPC/POPG lipids because of a lateral displacement of the TM4 by ∼1.5 Å (Cα r.m.s.d ∼1.24 Å, Extended Data Fig. 5i). The Ca^2+^ bound closed conformations in MSP1E3 DOPC/DOPG (PDBID: 6QMB) and MSP2N2 POPC/POPG are nearly identical (Cα r.m.s.d ∼0.50 Å). Finally, the intermediate conformation seen in the MSP2N2 DOPC/DOPG nanodiscs differs from the previously reported intermediate-closed one, as the TM4 is shifted further away from TM6 by ∼2 Å and accompanied with a counterclockwise rotation of the extracellular portion of TM4, such that there is minimal contact between TM4 and TM6 (Cα r.m.s.d ∼1.35 Å; Fig. 1lm and Extended Data Fig. 5j). This repositioning results in a conformation where the groove, while remaining isolated from the hydrocarbon core of the membrane, forms a continuous transmembrane pore that is wide enough to accommodate water molecules and permeant ions such as K^+^ and Cl^-^ (Fig. 1n-o). This is consistent with the reported channel activity of nhTMEM16 ^36,37^ ^38^. Therefore, we refer to this conformation as intermediate-open. Together, our results show that the scaffold protein and lipid composition affect both the distribution and structures of the conformations adopted by nhTMEM16.

### Basis of membrane thinning at the closed groove of nhTMEM16

The cryoEM map of Ca^2+^-bound nhTMEM16 with a closed groove in MSP1E3 nanodiscs reached an average resolution of 2.64 Å, allowing us to identify 36 well-defined non-protein densities, 18 on each protomer, that could be modeled as lipids (Fig. 2a-d and Extended Data Fig. 6). These densities define nearly continuous inner (IL) and outer leaflets (OL) and thus outline the protein-membrane interface (Fig. 2a-d and Extended Data Fig. 6a-e; Supplementary Video 1). In all cases the headgroups of lipids are not well resolved, therefore lipids were only built up to the phosphate atom. The 8 lipids near the dimer interface (D1-D8) define the inner and outer leaflets planes and adopt similar positions to those seen in Ca^2+^-bound open afTMEM16 ^30^ (Extended Data Fig. 6c-e), consistent with the idea this region undergoes minimal rearrangements during groove opening.

**Figure 2.**
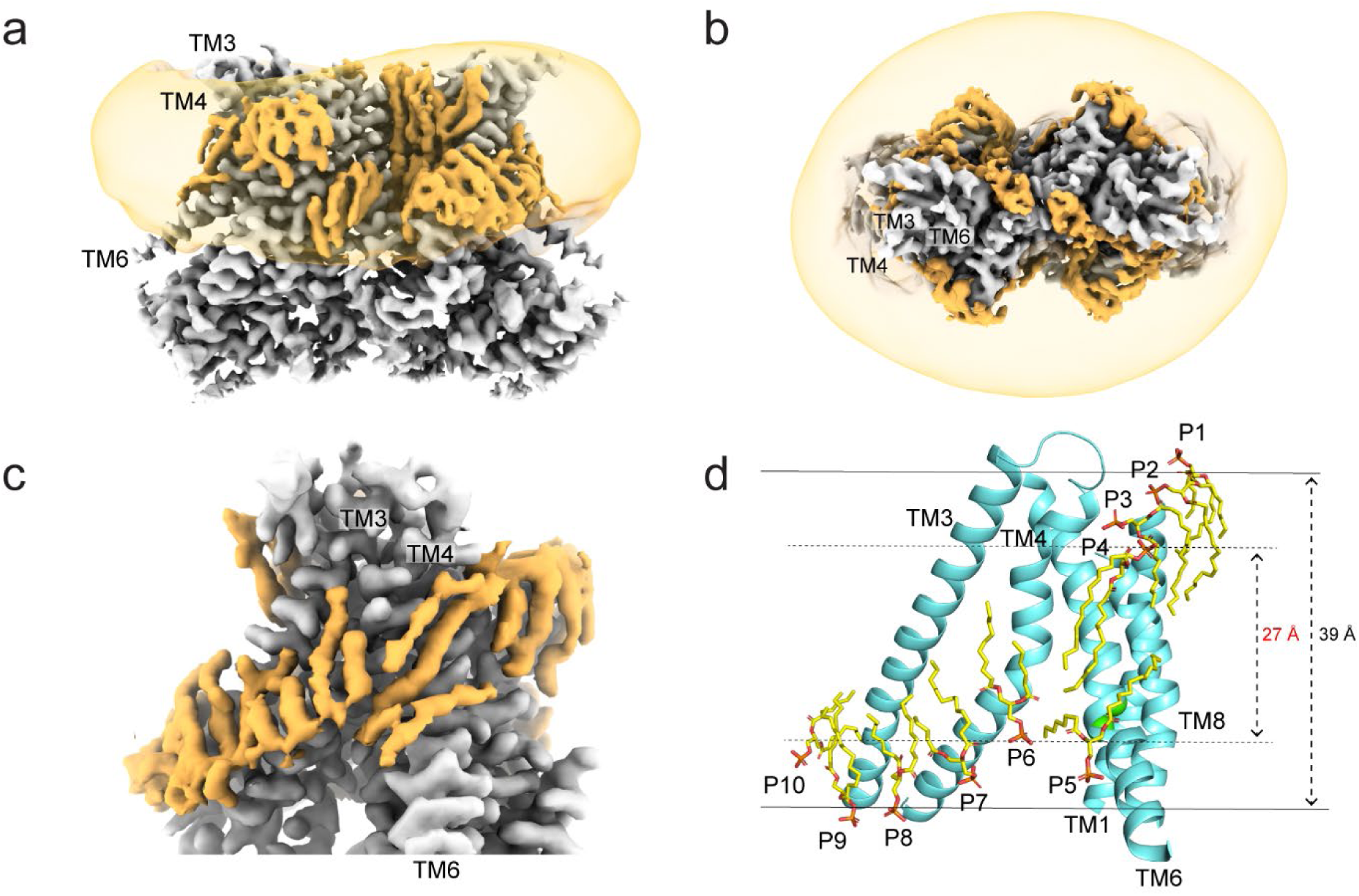
Arrangement of lipids at the closed groove of nhTMEM16. **a**-**b,** Segmented Cryo-EM map of nhTMEM16 in the Ca^2+^-bound closed state (gray), in MSP1E3 nanodiscs and DOPC/DOPG lipids, and the associated lipids (orange) viewed from the membrane plane (**a**) and from the extracellular side (**b**). The map showing the density of the nanodisc membrane is low-pass filtered to 10 Å and shown in transparent orange. **c,** View of the lipids outside of the closed groove from the plane of the membrane. **d,** Stick representation of the ten pathway lipids colored in yellow (P1-P10). Dashed arrow indicates the distance between the phosphate atoms of the last lipid from the inner (P6) and outer (P4) leaflets (∼27 Å) and the measured membrane thickness (∼39 Å). Lipids were built up to the phosphate atom in the head. Ca^2+^ ions are displayed as green spheres.

The arrangement of the 10 lipids near the pathway (P1-P10) reveals the structural basis of membrane thinning at a closed groove (Fig. 2d and Extended Data Fig. 6b). The outer leaflet P1-P4 lipids become progressively more tilted as they approach the site of TM4-TM6 contact that defines the groove closure (Fig. 2d) and their headgroups form extensive interactions with aromatic and polar residues in TM1, TM4, TM6, and TM8 (Fig. 3c). Notably, the headgroups of the P3 and P4 lipids adopt severely distorted poses (Fig. 2d) and interact with three aromatic residues, Y327 and F330 on TM4 and Y439 on TM6 (Fig. 3b). The P5 and P6 lipids delimit the inner leaflet across the wide intracellular vestibule of the pathway (Fig. 2d). The closest point of approach of the two leaflets at the closed groove is defined by the phosphate atoms of P4 and P6 which are separated by only ∼27 Å, so that the hydrocarbon core of the membrane is thinned by ∼30% (Fig. 2d). The Ca^2+^-bound and Ca^2+^-free closed conformations are nearly identical in this region (Extended Data Fig. 7a-e), indicating that protein-lipid interactions are likely unchanged in the two conformations. Thus, if the observed thinning is important for closed groove scrambling, then we expect these residues to play similar roles in the Ca^2+^-free and Ca^2+^-bound closed states.

**Figure 3.**
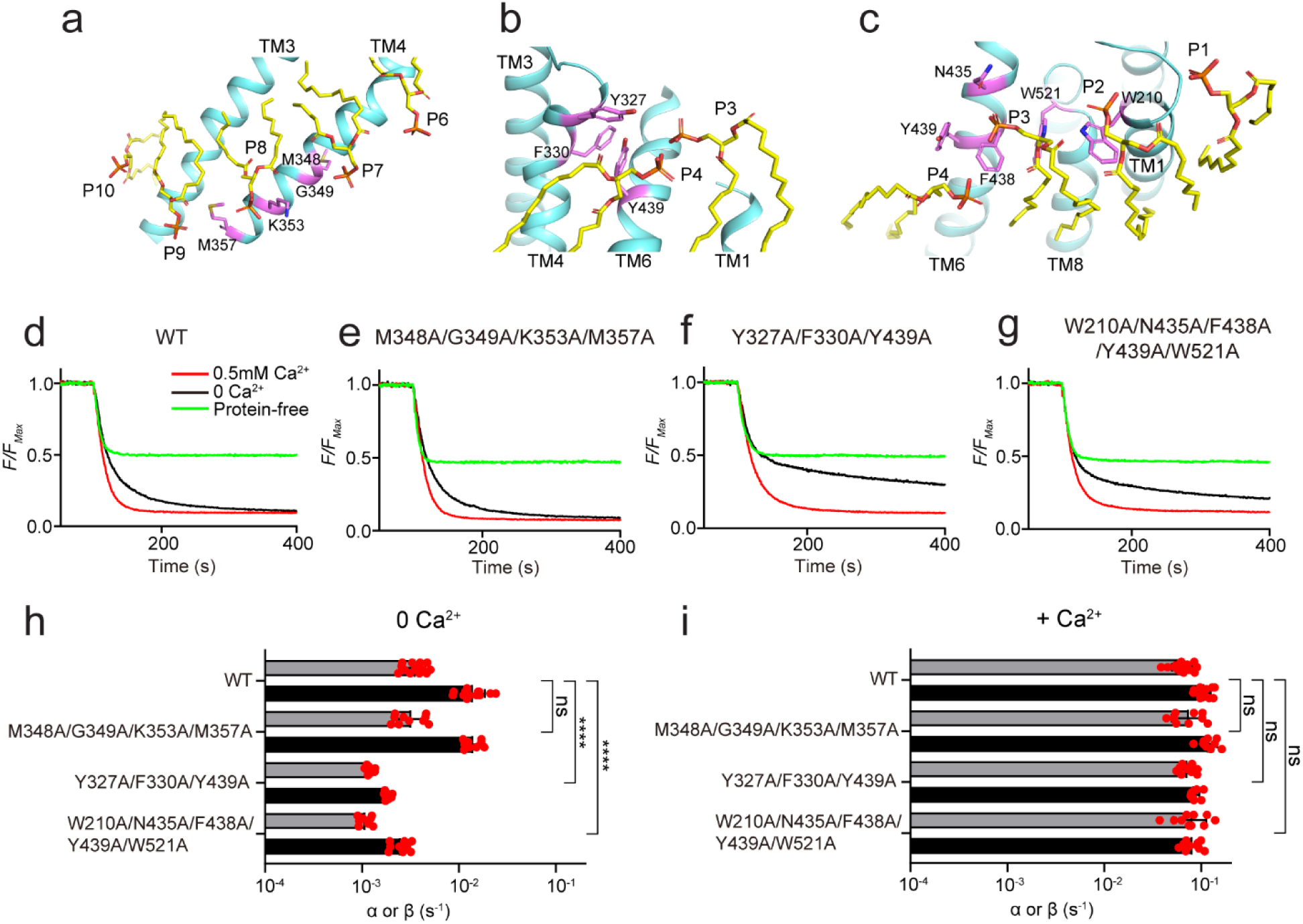
Identification of residues important for the closed groove scrambling. **a**-**c**, Residues coordinating the P7-P9 (**A**), P3-P4 (**B**) and P2-P4 lipids (**C**) in the Ca^2+^-bound closed state. **d**-**h**, Representative traces of the dithionite induced fluorescence decay in the scrambling assay for protein free liposomes (green), or WT and the mutants in the presence (red) and absence (black) of Ca^2+^. **h**-**i**, Forward (α) and reverse (β) scrambling rate constants of the mutants of nhTMEM16 aimed at disrupting the protein-lipid interactions (shown in **a**-**c**) measured in 0 (**h**) or 0.5 mM Ca^2+^ (**i**). Bars are averages for α (black) and β (gray) (N ≥ 3), error bars are S. Dev., and red circles are values from individual repeats. The statistical significance of the effects of the mutants on the scrambling rate constants was evaluated with a two-sided Student’s t-test with a Bonferroni correction. **** denotes p<10^-5^ after Bonferroni correction.

### Residues coordinating outer leaflet lipids are important in closed groove scrambling

To test this hypothesis, we designed three composite mutations aimed at disrupting coordination of the OL P1-P2-P3 (W210A/N435A/F438A/Y439A/W521A) and P3-P4 lipids (Y327A/F330A/Y439A) or of the IL P6-P10 lipids (M348A/G349A/M353A/K357A) (Fig. 3a-c) and tested their effects using a well-established in vitro assay to measure their effects on scrambling activity ^5,40,41^. In the presence of 0.5 mM Ca^2+^, when scrambling is primarily mediated by the open groove, the three constructs have WT-like rate constants (Fig. 3d-g, i). In contrast, in 0 Ca^2+^, when the groove is closed, the two constructs disrupting interactions with OL lipids display an 8- and 6-fold reduction in scrambling activity (p<10^-7^) (Fig. 3d-h). The scrambling rate constants of the mutant disrupting interactions with IL lipids are not statistically different from WT (P>0.4) (Fig. 3i). Thus, these mutants impair scrambling only in the absence of Ca^2+^, when the groove is closed, but have no effect in the presence of Ca^2+^, when the groove can open, suggesting that the two processes are mediated by different conformations. Therefore, we hypothesize that nhTMEM16, like afTMEM16 ^30^ and mTMEM16F ^19^, can scramble lipids with a closed groove and that remodeling of the OL lipids play a critical role in this process.

### Steps in groove opening of nhTMEM16

Our structures captured nhTMEM16 in several conformations, that likely reflect intermediates as the protein transitions from a Ca^2+^-free closed, to Ca^2+^-bound closed, intermediate, and open states (Fig. 4a-b). For simplicity, we consider only the conformations determined in MSP1E3 nanodiscs, as they have the highest resolution. In the Ca^2+^-free closed state the TM6 is in an α-helical configuration (Fig. 4a). Ca^2+^ binding induces a clockwise rotation of the intracellular portion of TM6 around A444, which results in the appearance of a π-helical turn that allows the E452 side chain to coordinate the bound Ca^2+^ ion in the upper and lower sites (Fig. 4a and Extended Data Fig. 7f, g). When the groove opens there is a counterclockwise rotation of the central portion of TM6 (between R432 and V447, Fig. 4b), which leads to the formation of a second π-helical turn (Fig. 4b). This disrupts the tight packing of TM4-TM6 in the closed groove and thus might facilitate opening (Fig. 4b). During these rearrangements the extracellular portion of TM6 does not move much, likely because the salt bridge between R432 on TM6 and E313 on TM3 provides a stabilizing interaction (Fig. 4a, b), suggesting it might play a role in the opening transition.

**Figure 4.**
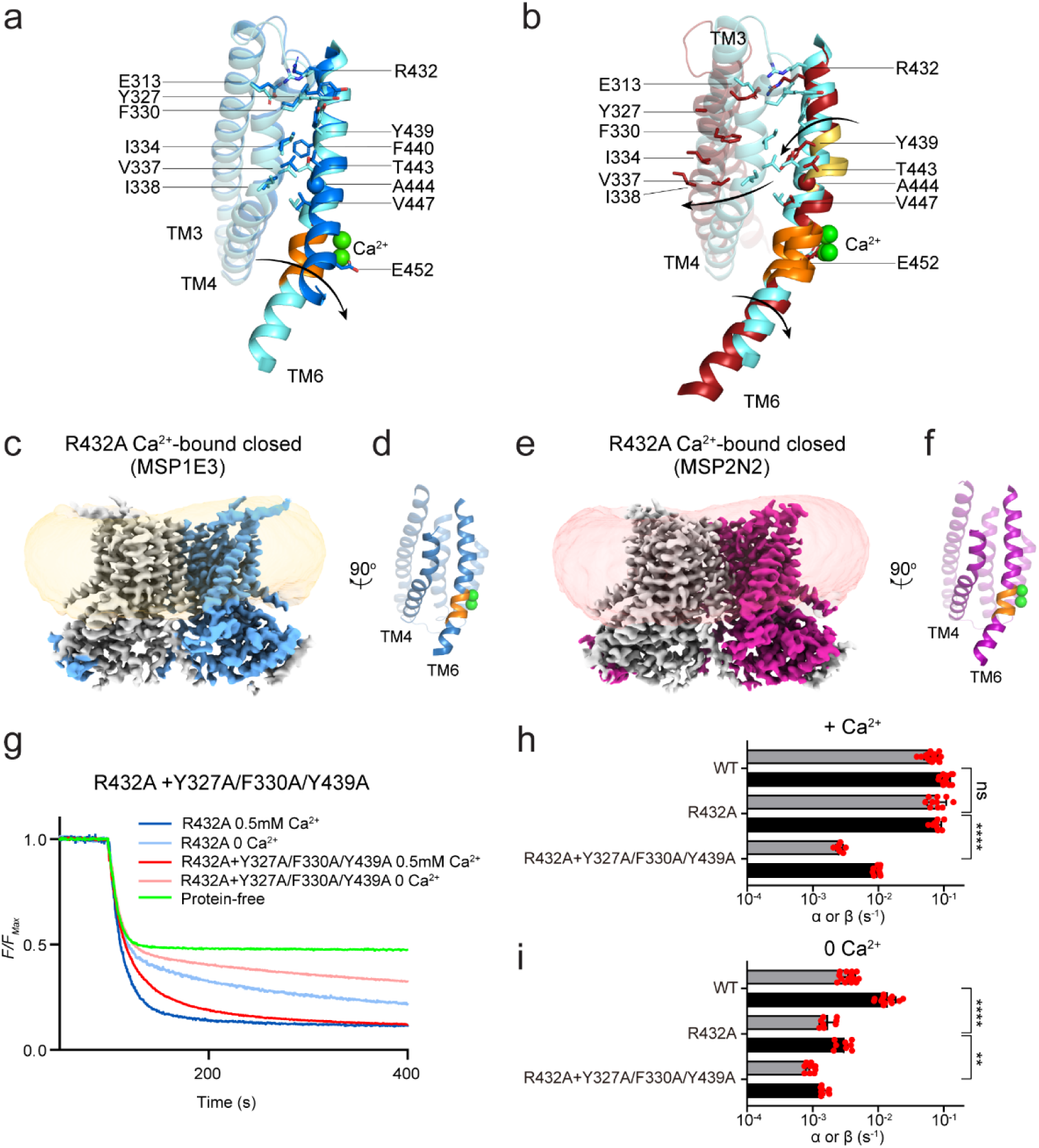
Role of the E313-R432 salt bridge in groove opening. **a**-**b**, Structural comparisons of the groove in apo (blue) vs Ca^2+^-bound closed (cyan) (**a**) and Ca^2+^-bound closed (cyan) vs open (red) states (**b**). Arrows denote rotations in the TM6. Sidechains of E313 and R432, and of the residues forming the TM4-TM6 interface are shown as sticks. The intra- and extra-cellular π-helical turns are colored in orange and gold, respectively. Ca^2+^ ions are displayed as green spheres. **c, d**, Cryo-EM maps of R423A nhTMEM16 in MSP1E3 (**c**) or MSP2N2 (**e**) nanodiscs. One protomer is colored in gray and the other in light blue (**c**) or magenta (**e**). The density of the nanodisc membrane is low pass filtered to 7 Å and shown in transparent orange (**c**) and red (**e**). **d, f**, Views of the groove from the plane of the membrane. TM4 and TM6 are shown in cartoon representation and labelled. Ca^2+^ ions are displayed as green spheres. **g**, Representative traces of the dithionite induced fluorescence decay in the scrambling assay for protein free liposomes (green), or R432A nhTMEM16 and the quadruple mutant R432A+Y327A/F330A/Y439A nhTMEM16 in the presence (dark blue and dark red) and absence (light blue and light red) of Ca^2+^. **h**-**i**, Forward (α) and reverse (β) scrambling rate constants of the mutants measured in 0.5 mM (**h**) or 0 Ca^2+^ (**i**). Bars are averages for α (black) and β (gray) (N ≥ 3), error bars are S. Dev., and red circles are values from individual repeats. The statistical significance of the effects of the mutants on the scrambling rate constants was evaluated with a two-sided Student’s t-test with a Bonferroni correction. ** denotes p<0.001 and **** denotes p< 10^-5^ after Bonferroni correction.

### Disruption of the E313-R432 salt bridge favors a closed groove

To test this hypothesis, we used cryoEM to image R432A nhTMEM16 in MSP1E3 or MSP2N2 nanodiscs in the presence of 0.5 mM Ca^2+^ (Fig. 4c, e and Extended Data Fig. 8a-g). The maps have similar average resolutions of 3.59 Å and 3.64 Å and show that R432A nhTMEM16 adopts a single class with a fully closed groove (Fig. 4d, f). Despite multiple rounds of iterative 3D classifications on dimers and protomers using different classification parameters and software (Extended Data Fig. 8e), no classes displaying an open groove could be identified. The models for Ca^2+^-bound R432A are nearly identical to that of WT Ca^2+^-bound closed nhTMEM16 in MSP1E3 nanodiscs (Fig. 2d and Extended Data Fig. 8h-j), with Cα r.m.s.d. of ∼0.93 and ∼0.98 Å. Consistently, the R432A map displays strong density in both Ca^2+^ binding sites (Extended Data Fig. 8k, l) and only the intracellular π-helical turn is present on TM6 (Fig. 4d, f). The only difference from the WT structure is that the density corresponding to the extracellular portion of the TM4 helix is weakened (Fig. 4d, f and Extended Data Fig. 8m, n), suggesting this region becomes dynamic in the mutant. Thus, disruption of the TM6-TM3 salt bridge favors a closed groove conformation.

Next, we determined how the R432A mutation affects scrambling activity in DOPC/DOPG liposomes and found it retains WT-like activity in the presence of Ca^2+^ and a ∼4-fold reduction in 0 Ca^2+^ (p<10^-5^) (Fig. 4g-i). As the only visible conformation of Ca^2+^-bound R432A nhTMEM16 in MSP1E3 and MSP2N2 nanodiscs is with a closed groove, we hypothesize its activity in the presence of Ca^2+^ reflects closed-groove scrambling. We tested this hypothesis by combining the R432A mutant with the triple Y327A/F330A/Y439A mutant which only impairs scrambling in 0 Ca^2+^ (Fig. 3d), when the groove is closed. We expected that if Ca^2+^ bound R432A mediates closed groove scrambling, then the quadruple mutant should display reduced activity in the presence of Ca^2+^, while neither of the original mutants did. In contrast, if scrambling by Ca^2+^-bound R432A occurs via an open groove mechanism, we would expect the quadruple mutant to have WT-like activity. Results meet expectations: the R432A+Y327A/F330A/Y439A mutant displays an ∼8-fold reduction in scrambling activity in 0.5 mM Ca^2+^ (p<10^-5^) (Fig. 4g-h), very similar to the effect of the triple Y327A/F330A/Y439A in 0 Ca^2+^ (Fig. 3d). In the absence of Ca^2+^ the quadruple mutant reduces scrambling activity ∼9-fold (Fig. 4g, i). Together, these results suggest that the R432A mutation favors a closed groove conformation by causing a decrease in Ca^2+^ potency and that nhTMEM16 mediates Ca^2+^-dependent closed-groove scrambling. However, we cannot rule out the possibility that the chemical step in the dithionite assay limits our ability to accurately resolve the effects of the mutants in the presence of Ca^2+^. Similarly, it is possible that the open probability of the Ca^2+^ bound R432A groove is too low to be detected in cryoEM imaging experiments but sufficient to account for the observed activity in liposomes. Finally, we note that the R432A mutation might also slightly perturb the conformation of the closed groove as it mildly impairs scrambling in 0 Ca^2+^ (Fig. 4i).

### Straightening of TM6 is important for closed-groove scrambling

One key assumption of the closed groove scrambling model is that the structural differences between the apo-closed and Ca^2+^-bound closed conformations underlie the Ca^2+^ dependence of some TMEM16 scramblases. In our structures the major difference occurring on Ca^2+^ binding is the straightening of the intracellular portion of the TM6 helix around A444 and the formation of the intracellular π-helical turn (Fig. 4a and Extended Data Fig. 7f, g). To investigate whether this rearrangement modulates closed groove scrambling we introduced a helix-bending proline at position A444. The A444P mutant displays severely impaired scrambling activity in DOPC/DOPG liposomes, with a >20-fold reduction in 0.5 mM Ca^2+^ and ∼17-fold reduction in 0 Ca^2+^ (p<10^-8^) (Fig. 5a-c). Cryo-EM imaging of A444P nhTMEM16 in MSP1E3 DOPC/DOPG nanodiscs in 0.5 mM Ca^2+^ (Fig. 5d-I and Extended Data Fig. 9a-i) shows the mutant exclusively adopts conformations with a closed groove (Cα r.m.s.d. <0.4 Å to WT Ca^2+^-bound nhTMEM16 in all cases). Extensive classification results in four dimer classes: three are symmetrical and differ only in the occupancy of the Ca^2+^ binding sites and in the conformation of the intracellular portion of TM6 below 444P (Fig. 5d, f and Extended Data Fig. 9j-l), and we refer to them as ‘long TM6’ (∼1.3% of particles), ‘short TM6’ (∼45% of particles), and ‘bent TM6’ (∼4.5% of particles). The fourth class is asymmetric, with one protomer in the long TM6 and one in the short TM6 conformations (21.8% of particles) (Extended Data Fig. 9a-i). In the remaining ∼27% of particles the density of the TM6 helices is not well defined, and therefore could not be assigned to any of the classes. Whereas in the long TM6 map the density for both Ca^2+^ binding sites is similar (Fig. 5d, g and Extended Data Fig. 9j), in the short TM6 map the density in the upper Ca^2+^ binding site is weaker than that in the bottom site (Fig. 5e, h and Extended Data Fig. 9k). In the bent TM6 conformation the E452 side chain on TM6 is oriented away from the Ca^2+^ binding site, and we detect weak density only for the bottom Ca^2+^ binding site (Fig. 5f, i and Extended Data Fig. 9l), suggesting impaired ion binding. Thus, the A444P mutation impairs scrambling, and favors states with short or bent TM6, with weakened Ca^2+^-binding. These results suggest that the Ca^2+^-dependent straightening of TM6 might be important for closed groove scrambling. It is likely that the contribution of the TM6 rearrangements to scrambling are smaller than those of groove opening. Therefore, their effects are revealed only in conditions favoring closed groove scrambling.

**Figure 5.**
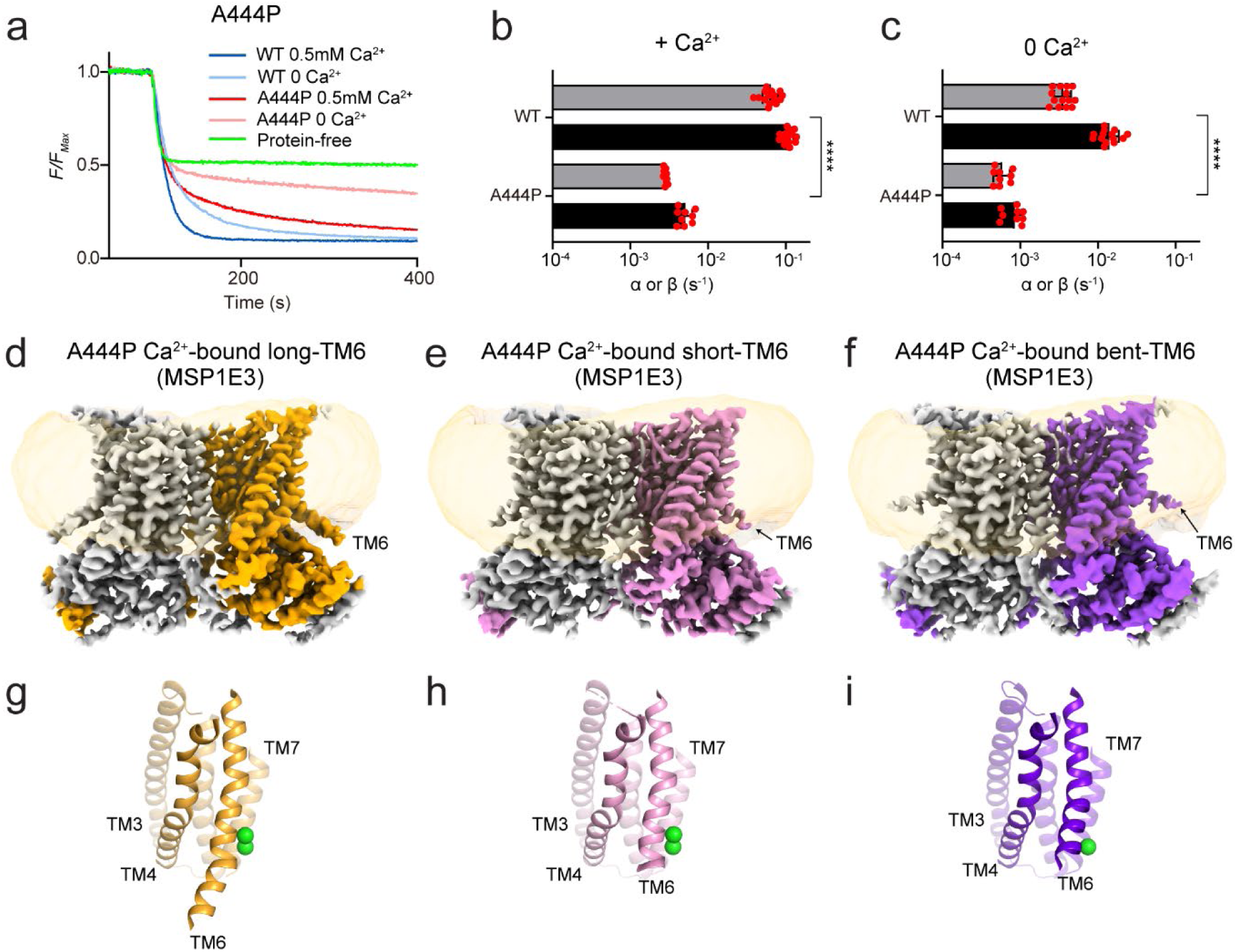
Disruption of TM6 straightening impairs lipid scrambling. **a**, Representative traces of the dithionite induced fluorescence decay in the scrambling assay in protein-free liposomes (green) or proteoliposomes reconstituted with WT and A444P nhTMEM16 in the presence (dark blue and dark red) and absence (light blue and light red) of Ca^2+^. **b**-**c**, Forward (α) and reverse (β) scrambling rate constants of the mutants measured in 0.5 mM (**b**) or 0 Ca^2+^ (**c**). Bars are averages for α (black) and β (gray) (N ≥ 3), error bars are S. Dev., and red circles are values from individual repeats. The statistical significance of the effects of the mutants on the scrambling rate constants was evaluated with a two-sided Student’s t-test with a Bonferroni correction. **** denotes p<10^-5^ after Bonferroni correction. **d**-**f**, Cryo-EM maps of A444P nhTMEM16 in MSP1E3 nanodiscs in the long TM6 state (**d**), short TM6 state (**e**) and bent TM6 state (**f**). One protomer is colored in gray and the other in orange (**d**), pink (**e**), and purple (**f**). The density of the nanodisc membrane is low pass filtered to 7 Å and shown in transparent orange. **g**-**i**, The groove in the long TM6 (**g**), short TM6 (**h**) and bent TM6 (**i**) states is viewed from the plane of the membrane. Transmembrane helices are labelled and Ca^2+^ ions are displayed as green spheres.

### The effect of nhTMEM16 mutants depends on the lipid environment

Finally, we note that our results on R432A nhTMEM16 contrast with our previous report ^27^, where we showed that this and several other mutations, such as Y439A, greatly impair nhTMEm16 scrambling activity. The key difference between our current and past experiments is the membrane composition, a DOPC/DOPG mixture here and a POPE/POPG/Egg PC mixture then. To test if changes in lipid headgroups or acyl chain saturation underlie this difference we reconstituted WT and mutant nhTMEM16 in liposomes formed from DOPC/DOPG, POPE/POPG, DOPE/DOPG and POPC/POPG lipid mixtures. Consistent with previous reports ^27,37^, WT nhTMEM16 is active in all compositions, with a ∼3-fold reduction in POPE/POPG membranes compared to the other conditions (Fig. 6a, d and e). In contrast, scrambling by R432A nhTMEM16 is reduced by >200-fold in POPE/POPG liposomes, but is WT-like in DOPC/DOPG, DOPE/DOPG and POPC/POPG membranes (Fig. 6b, d and e). The Y439A mutant is also only inhibited in POPE/POPG liposomes and WT-like in the others (Fig. 6c-e). Thus, scrambling activity is specifically reduced in POPE-containing membranes, as all constructs are functional in DO lipids as well as in PO membranes formed from PC/PG mixtures. This rules out specific interactions between the protein and the PE headgroup or the 16:1-18:0 acyl chains of PO lipids. While we do not have a satisfactory explanation for this observation, we note that the transition temperature (T_m_) of POPE membranes is close to room temperature, whereas those of all other tested lipids is < 0 °C ^42^. This raises the possibility that scrambling might be sensitive to the formation of local gel-like microdomains with reduced fluidity. This finding underscores the importance of the membrane environment in assaying the activity of TMEM16 scramblases.

**Figure 6.**
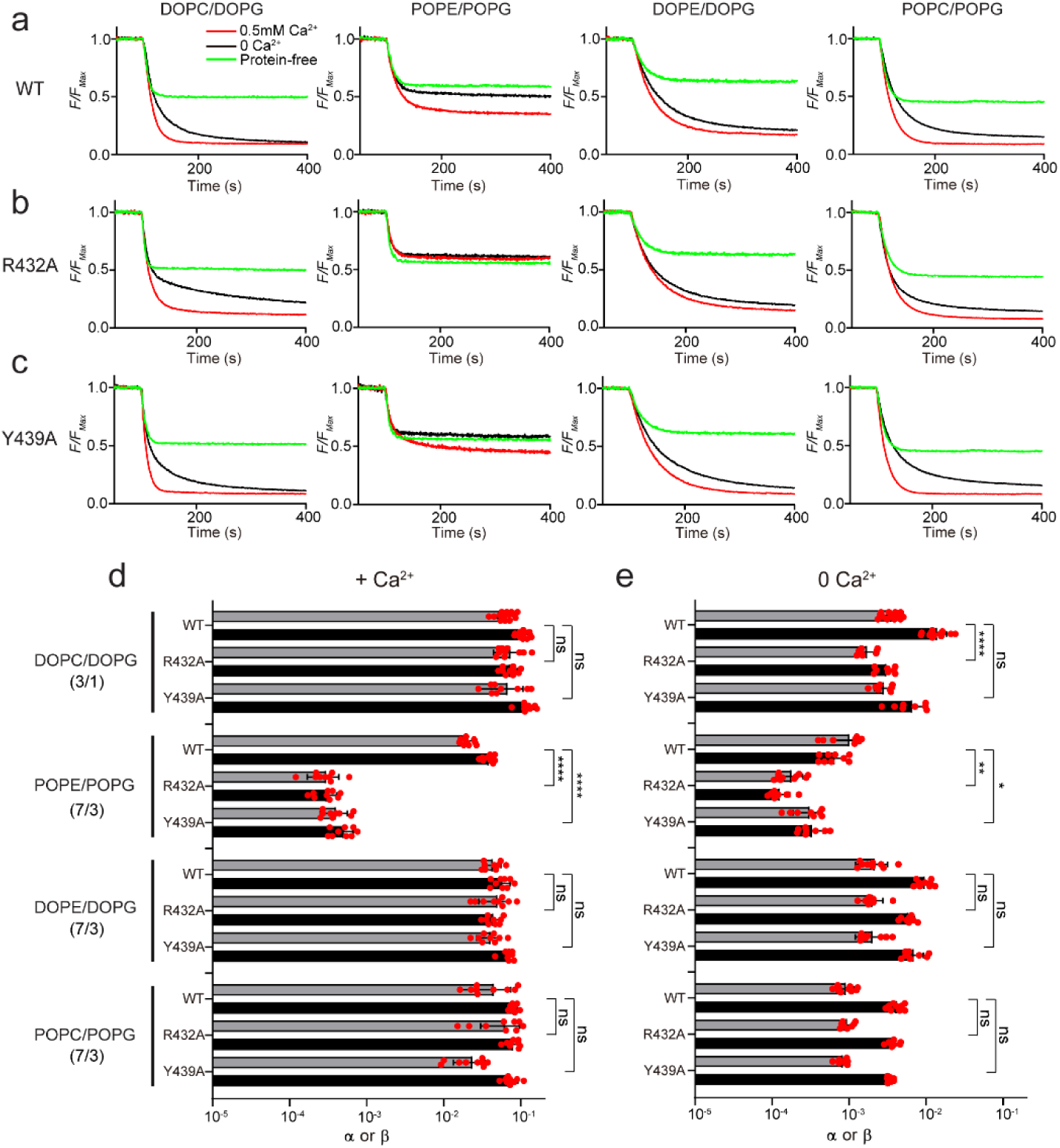
The lipid environment determines the effect of mutations of nhTMEM16. **a-c,** Representative traces of the dithionite induced fluorescence decay in the scrambling assay in protein-free liposomes (green) or proteoliposomes reconstituted with WT (**a**), R432A (**b**) or Y432A (**c**) nhTMEM16 in the presence (red) and absence (black) of Ca^2+^. Liposomes were formed from a 3:1 mix of DOPC/DOPG, or a 7:3 mix of POPE/POPG, DOPE/DOPG and POPC/POPG lipids. **d-e,** Forward (α) and reverse (β) scrambling rate constants of WT, R423A and Y439A nhTMEM16 in the four different lipid compositions (as in **a**-**c**) in the presence of 0.5 mM (**d**) or 0 Ca^2+^ (**e**). Bars are averages for α (black) and β (gray) (N ≥ 3), error bars are S. Dev., and red circles are values from individual repeats. The statistical significance of the effects of the mutants on the scrambling rate constants was evaluated with a two-sided Student’s t-test with a Bonferroni correction. *, **, and **** respectively denote p<5.10^-3^, <10^-3^, and <10^-5^ after Bonferroni correction.

## Discussion

The Ca^2+^ dependent activation of TMEM16 scramblases plays a critical role in a variety of cellular signaling pathways ^1-3,6^. Despite extensive structural and functional investigations, the structural basis of their Ca^2+^-dependent activation and lipid transport mechanisms remain poorly understood. For example, TMEM16 activation is thought to entail opening of a hydrophilic groove following Ca^2+^ binding to a site formed by residues in the TM6-TM8 helices. However, it is not clear how Ca^2+^ binding facilitates the displacement of TM4 to open the groove. Further, while an open groove facilitates scrambling, the closed groove has also been proposed to be scrambling competent ^19,30^. However, the structural basis of lipid scrambling outside a closed groove remain unknown.

Our high-resolution structures of Ca^2+^-bound closed nhTMEM16 reconstituted into lipid nanodiscs allowed us to directly visualize how lipid molecules interact with the closed groove of the scramblase (Fig. 2a-d; Supplementary Video 1 and Extended Data Fig. 6). We found that inner and outer leaflet lipids adopt distorted poses where their phosphate headgroups coming within ∼27 Å of each other at the closed groove. This arrangement of the lipids results in a ∼30% thinning of the hydrocarbon core of the membrane which will facilitate scrambling when the groove is closed. Mutants aimed at disrupting this lipid arrangement only impair scrambling by nhTMEM16 in the absence of Ca^2+^, when the groove is closed, but have no effect in the presence of Ca^2+^, when the groove can open (Fig. 3h, i). These observations suggest that the observed distortion of the lipids near the groove, and the ensuing membrane thinning, are critical for closed groove scrambling. We speculate this mechanism is likely conserved across TMEM16 scramblases for several reasons. First, mutations at similar positions also impair closed groove scrambling in afTMEM16 ^30^. Second, the membrane near the closed groove of afTMEM16 and TMEM16F is thinned to a similar extent to what we see here for nhTMEM16 ^19,21,30^ and, third, in TMEM16K two resolved detergent molecules adopt tilted poses similar to those of the P4 and P8 lipids in nhTMEM16 ^18^ (Extended Data Fig. 10a-c). These results suggest that membrane thinning could be a general and evolutionary conserved mechanism for lipid scrambling by the TMEM16s, although with different details among homologues. With a closed groove, thinning is less pronounced, and scrambling is slower than when the groove is open, rationalizing the Ca^2+^ dependence of TMEM16 activity (Extended Data Fig. 10d-f). The strong dependence of closed groove scrambling on membrane properties could provide a mode of regulation of TMEM16 activity in cellular membranes, such as the cholesterol rich plasma membrane or the thinner ER membrane. In sum, our work ^17,30^, together with that of others ^19,21,22,29,31^, suggest that the TMEM16 proteins provide a delocalized, low-energy path for lipids to traverse the membrane, rather than forming discrete binding sites that the lipids move between. Mutations aimed at disrupting protein-lipid interactions near an open or closed groove leave scrambling unaltered ^30^ or cause less than a tenfold reduction in activity (Fig. 3), respectively. Thus, while the membrane distortion imposed by these protein-lipid interactions is critical for scrambling, it is possible that the transported lipids might not be the ones visualized to directly interact with the protein.

Piecing together our results with past work ^7,21^ we can provide a detailed description of the molecular transitions underlying the Ca^2+^ dependent activation of nhTMEM16 (Supplementary Video 2). To this end, we make two plausible assumptions: that the previously reported state obtained in Ca^2+^-free conditions corresponds to a partially bound state and, that the 7 known conformations of nhTMEM16, reported here and earlier ^21^, represent intermediates along the transition from apo closed to Ca^2+^ bound open. The relative stability of these conformations might be differentially affected by the membrane environment, which would result in slightly different opening trajectories. Nonetheless, we hypothesize that these states are visited -albeit briefly-during opening. Starting in the apo conformation the groove is closed, TM6 is bent, and the cytosolic domains are apart (PDBID: 8TPM) (Fig. 1a, f). Partial occupancy of the Ca^2+^ sites promotes a movement of the cytosolic domains away from the symmetry axis and a partial straightening of TM6 (PDBID: 6QM4) (Extended Data Fig. 5a, e and g). Binding of both Ca^2+^ ions favors the compete straightening of TM6 and the formation of the intracellular π-helix turn, which allows the acidic side chain of E452 on TM6 to come within coordination distance of the bound ion (PDBID: 8TOI) (Fig. 1b, g, 4a and Extended Data Fig. 7f). These rearrangements weaken the TM4-TM6 packing, allowing TM4 to move and adopt the intermediate-closed (PDBID: 6QMA) and intermediate-open configurations (PDBID: 8TPN) (Fig. 1d, i), which might lead to the formation of ion conduction pores (Fig. 1n, o). The partial disengagement of the TM4 from TM6 allows the extracellular portion of TM6 between the bound Ca^2+^ ions and the R432-E313 salt bridge to rotate, resulting in the formation of a second π-helix turn at A444 (Fig. 4b). These rearrangements disrupt the TM4-TM6 interface, and likely facilitate the transition to conformations with an open groove (PDBID: 8TOL and 8TOK) (Fig. 1c, e, h and j). The role of the R432-E313 salt bridge in groove opening is supported by our finding that the R432A mutation stabilizes the Ca^2+^-bound closed conformation in nhTMEM16 (Fig. 4c-f). This interaction between TM6 and TM3 is well conserved across fungal and mammalian TMEM16 homologues (Extended Data Fig. 10g-j). Interestingly, in mTMEM16F the interaction is present, but the charges are reversed, and the closed groove is the most favored conformation ^19,22^. Similarly, charge-reversal mutants of nhTMEM16 are non-functional in POPE-containing liposomes ^27^, suggesting that the precise arrangement of charges in the salt bridge is also important.

Our results reveal that the environment plays a critical role in determining the structures of nhTMEM16 conformations as well as the population distribution among them. Indeed, subtle changes in lipid composition, from PO to DO lipids, and/or nanodisc scaffold proteins, from MSP1E3 to MSP2N2, can affect ion binding, protein conformations, and their distributions. Specifically, in MSP1E3 DOPC/DOPG nanodiscs nhTMEM16 can adopt an apo conformation (Fig. 1a, f and Extended Data Fig. 5d) which differs from the previously reported one in MSP2N2 POPC/POPG nanodiscs because of conformational rearrangements in the cytosolic and groove regions (Extended Data Fig. 5c). These differences likely reflect the different occupancy of the Ca^2+^ sites, empty in our structure but partially occupied by an unknown ligand in the previous ^21^ one (Extended Data Fig. 5e). This suggests that the membrane environment might modulate TMEM16 activity by affecting ion binding, as both structures were obtained in similar Ca^2+^ buffering conditions. Structural differences are also observed in the conformations adopted in the MSP2N2 DOPC/DOPG nanodiscs compared to those seen in MSP2N2 POPC/POPG ^21^, as both the open and intermediate groove conformations in DO lipids are more open than those seen in PO lipids (Fig. 1l, m and Extended Data Fig. 5i, j). Indeed, in the intermediate-open conformation seen in DO lipids TM4 makes minimal contact with TM6 (Fig. 1n) leading to the formation of a continuous transmembrane pore with internal diameter >4.1 Å, which is sufficiently wide to be fully hydrated and to allow permeation of both chloride and potassium ions. We hypothesize this state could correspond to an ion conductive conformation of nhTMEM16. Unexpectedly, the conformations seen in the smaller MSP1E3 DOPC/DOPG nanodiscs are indistinguishable from those reported in MSP2N2 POPC/POPG, indicating that both nanodisc diameter and lipid properties contribute to stabilizing specific conformations of the protein. Strikingly, these differences in particle distributions, ligand binding state, and conformations do not correlate with changes in activity, as nhTMEM16 scrambles lipids equally well in DOPC/DOPG and POPC/POPG liposomes (Fig. 6), the Ca^2+^-free conditions are nominally identical (Fig. 1a) ^21^, and the lipid composition is the same in our MSP1E3 and MSP2N2 reconstitutions (Fig. 1k). Finally, our results suggest that reconstitution in the smaller MSP1E3 nanodiscs favors closed or intermediate conformations of nhTMEM16, as we observe a ∼20-fold increase in the fraction of particles with an open groove in the larger MSP2N2 discs (Fig. 1k), where the fraction of particles with a closed groove is undetectable (Fig. 1k). This raises the possibility that unbounded membranes, such as those of liposomes, might perturb less the conformational landscape of the imaged proteins.

Recently, much work has been devoted to extracting free energy landscapes of proteins from the analysis of particle distributions in cryoEM imaging experiments ^43–48^. Our findings challenge the straightforward assumption that the particle distribution is a faithful reporter of the conformational landscape of the protein. The lack of correspondence between protein activity, particle distribution, and conformation confounds the correlation between the free-energy landscape of a protein, cryoEM particle counts, and activity. Rather, our results show that these distributions also reflect the contributions of the environment, and that even subtle changes in detergent, scaffold protein, or lipid constituents of nanodiscs can shift the preferred conformations adopted by the reconstituted protein. Our results suggest that the energetic contributions of these environmental factors could be relatively small, in the 1-3 kCal Mol^-1^ range. Nonetheless, their effects can profoundly affect the interpretation of structural experiments such as the identification of dominant conformations in a sample (Fig. 1k) or in the assignment of functional properties to a specific conformation (Fig. 1o). These effects should be evaluated for each protein system under consideration, as for example whereas the conformational landscape of WT nhTMEM16 is highly sensitive to the environment, that of its close homologues afTMEM16 or mTMEM16F is not ^17,19,22,30^, and even the single point mutant R432A nhTMEM16 exclusively adopts only one conformation in different nanodisc scaffolds (Fig. 4c-f). Interestingly, a recent report suggests that the conformational landscape of the pentameric ligand gated ion channel ELIC is also affected by subtle changes in the imaging environment ^49^, suggesting these considerations are generally applicable to diverse types of membrane proteins. It will be interesting to see whether structures determined in unbounded membranes, such as liposomes will differ from those determined in nanodiscs or detergent systems.

## Data availability

The data that support this study are available from the corresponding author upon request. All models and associated cryoEM maps have been deposited into the Electron Microscopy Data Bank (EMDB) and the Protein Data Bank (PDB). The depositions include final maps, unsharpened maps, local refined maps, and associated FSC curves. Accession codes are listed here and in Table 1. WT nhTMEM16 in MSP1E3 nanodiscs in the absence of Ca^2+^: EMD-41477 and PDB 8TPM; WT nhTMEM16 in MSP1E3 nanodiscs in the presence of Ca^2+^ (closed state): EMD-41453, EMD-41457 (consensus map), EMD-41458 (monomer map) and PDB 8TOI; WT nhTMEM16 in MSP1E3 nanodiscs in the presence of Ca^2+^ (open state): EMD-41455 and PDB 8TOL; WT nhTMEM16 in MSP2N2 nanodiscs in the presence of Ca^2+^ (intermediate-open state): EMD-41478 and PDB 8TPN; WT nhTMEM16 in MSP2N2 nanodiscs in the presence of Ca^2+^ (open state): EMD-41454 and PDB 8TOK; R432A nhTMEM16 in MSP1E3 nanodiscs in the presence of Ca^2+^ (closed state): EMD-41479 and PDB 8TPO; R423A nhTMEM16 in MSP2N2 nanodiscs in the presence of Ca^2+^ (closed state): EMD-41480 and PDB 8TPP; A444P nhTMEM16 in MSP1E3 nanodiscs in the presence of Ca^2+^: EMD-41481 and PDB 8TPQ for the closed state with long TM6, EMD-41482 and PDB 8TPR for the closed state with short TM6, EMD-41483 and PDB 8TPS for the closed state with bent TM6, EMD-41484 and PDB 8TPT for the closed state with asymmetric TM6 (long TM6/short TM6).

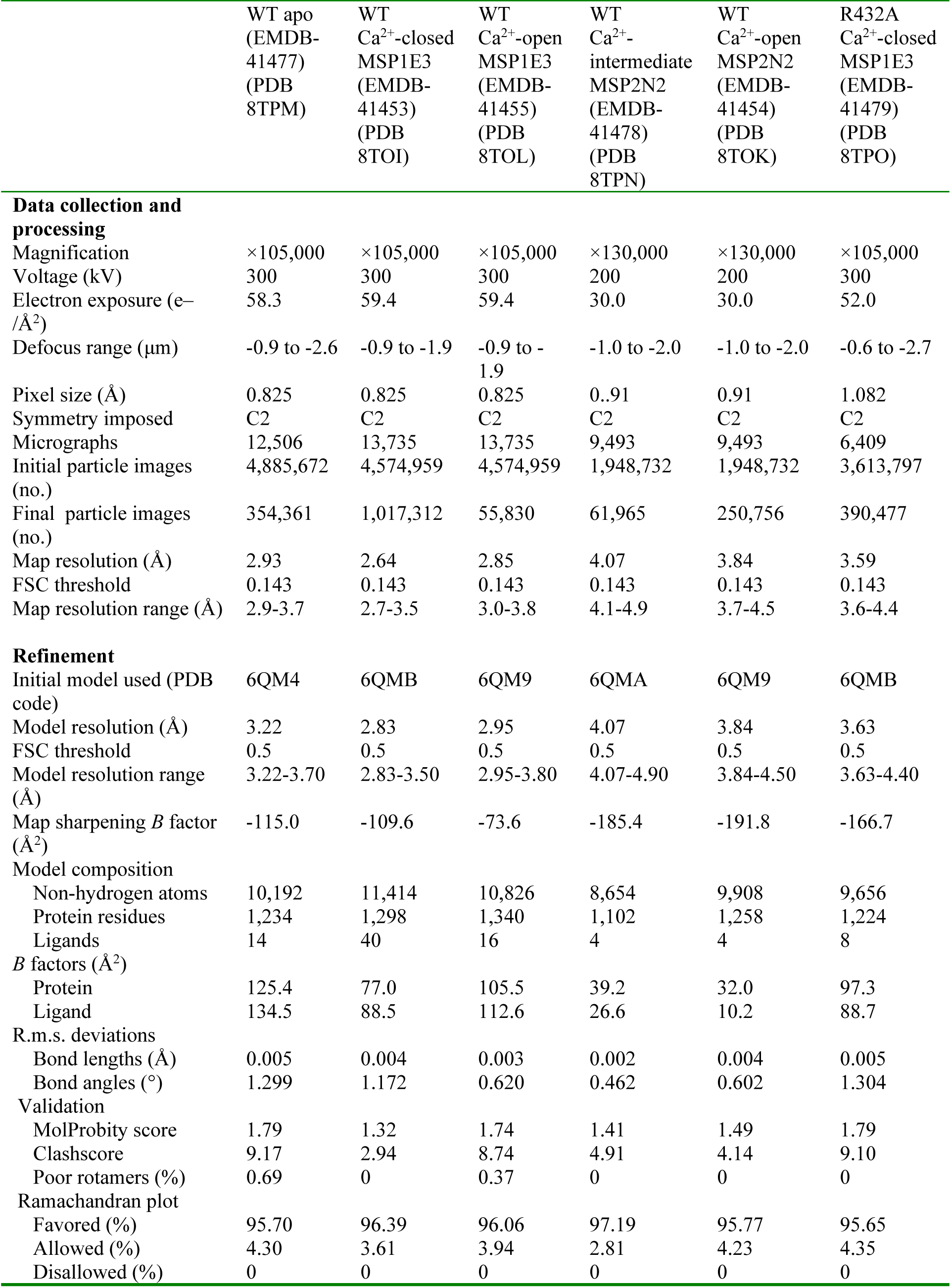
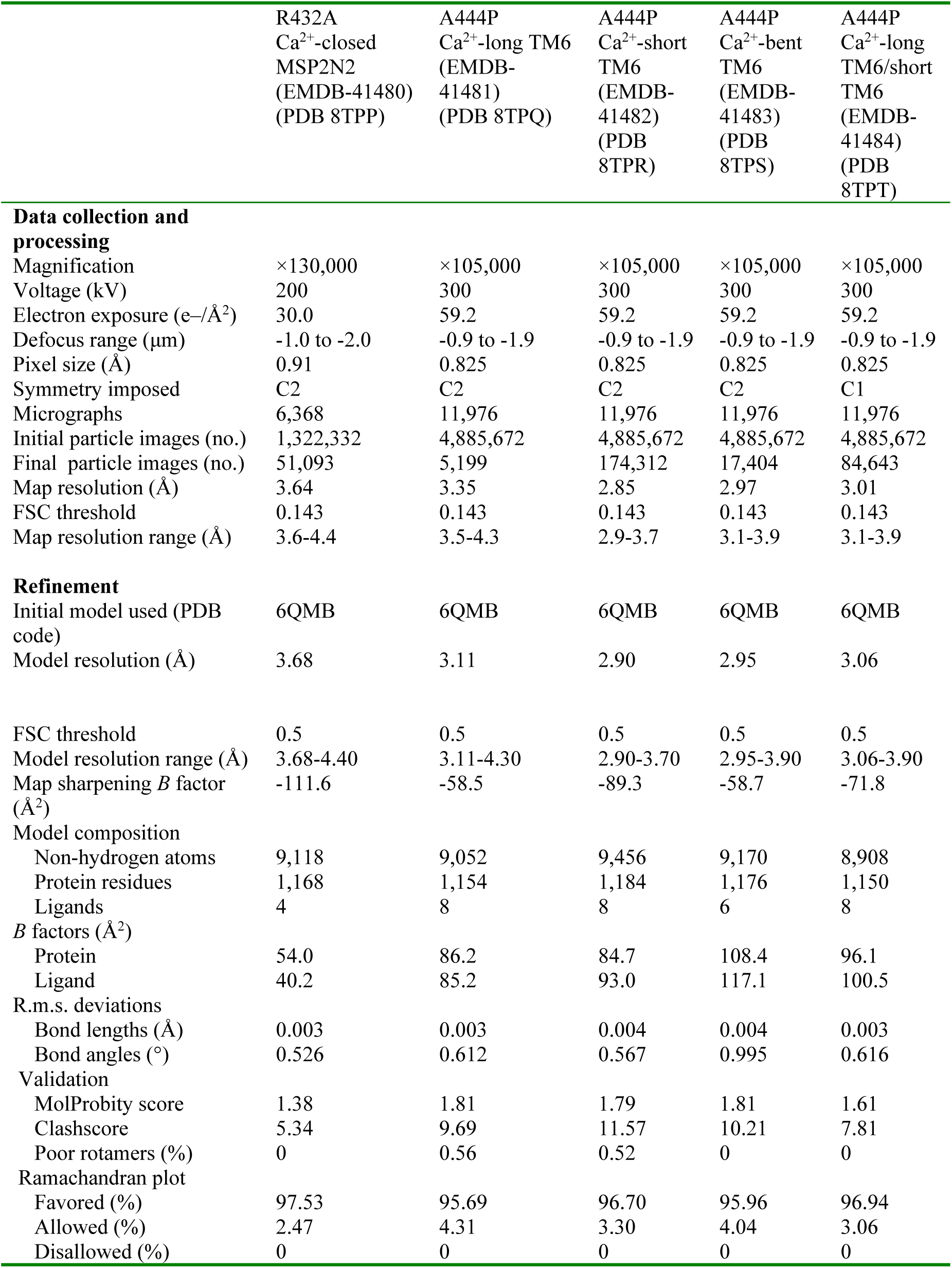
Cryo-EM data collection, refinement and validation statistics.

Cryo-EM data collection, refinement and validation statistics
|  | WT apo<br>(EMDB-<br>41477)<br>(PDB<br>8TPM) | WT<br>Ca<sup>2+</sup>-closed<br>MSP1E3<br>(EMDB-<br>41453)<br>(PDB<br>8TOI) | WT<br>Ca<sup>2+</sup>-open<br>MSP1E3<br>(EMDB-<br>41455)<br>(PDB<br>8TOL) | WT<br>Ca<sup>2+</sup>-<br>intermediate<br>MSP2N2<br>(EMDB-<br>41478)<br>(PDB<br>8TPN) | WT<br>Ca<sup>2+</sup>-open<br>MSP2N2<br>(EMDB-<br>41454)<br>(PDB<br>8TOK) | R432A<br>Ca<sup>2+</sup>-closed<br>MSP1E3<br>(EMDB-<br>41479)<br>(PDB<br>8TPO) |
| --- | --- | --- | --- | --- | --- | --- |
| **Data collection and processing** |  |  |  |  |  |  |
| Magnification | ×105,000 | ×105,000 | ×105,000 | ×130,000 | ×130,000 | ×105,000 |
| Voltage (kV) | 300 | 300 | 300 | 200 | 200 | 300 |
| Electron exposure (e<sup>-</sup>/Å<sup>2</sup>) | 58.3 | 59.4 | 59.4 | 30.0 | 30.0 | 52.0 |
| Defocus range (μm) | -0.9 to -2.6 | -0.9 to -1.9 | -0.9 to -1.9 | -1.0 to -2.0 | -1.0 to -2.0 | -0.6 to -2.7 |
| Pixel size (Å) | 0.825 | 0.825 | 0.825 | 0.91 | 0.91 | 1.082 |
| Symmetry imposed | C2 | C2 | C2 | C2 | C2 | C2 |
| Micrographs | 12,506 | 13,735 | 13,735 | 9,493 | 9,493 | 6,409 |
| Initial particle images (no.) | 4,885,672 | 4,574,959 | 4,574,959 | 1,948,732 | 1,948,732 | 3,613,797 |
| Final particle images (no.) | 354,361 | 1,017,312 | 55,830 | 61,965 | 250,756 | 390,477 |
| Map resolution (Å) | 2.93 | 2.64 | 2.85 | 4.07 | 3.84 | 3.59 |
| FSC threshold | 0.143 | 0.143 | 0.143 | 0.143 | 0.143 | 0.143 |
| Map resolution range (Å) | 2.9-3.7 | 2.7-3.5 | 3.0-3.8 | 4.1-4.9 | 3.7-4.5 | 3.6-4.4 |
| **Refinement** |  |  |  |  |  |  |
| Initial model used (PDB code) | 6QM4 | 6QMB | 6QM9 | 6QMA | 6QM9 | 6QMB |
| Model resolution (Å) | 3.22 | 2.83 | 2.95 | 4.07 | 3.84 | 3.63 |
| FSC threshold | 0.5 | 0.5 | 0.5 | 0.5 | 0.5 | 0.5 |
| Model resolution range (Å) | 3.22-3.70 | 2.83-3.50 | 2.95-3.80 | 4.07-4.90 | 3.84-4.50 | 3.63-4.40 |
| Map sharpening *B* factor (Å<sup>2</sup>) | -115.0 | -109.6 | -73.6 | -185.4 | -191.8 | -166.7 |
| **Model composition** |  |  |  |  |  |  |
| Non-hydrogen atoms | 10,192 | 11,414 | 10,826 | 8,654 | 9,908 | 9,656 |
| Protein residues | 1,234 | 1,298 | 1,340 | 1,102 | 1,258 | 1,224 |
| Ligands | 14 | 40 | 16 | 4 | 4 | 8 |
| ***B* factors (Å<sup>2</sup>)** |  |  |  |  |  |  |
| Protein | 125.4 | 77.0 | 105.5 | 39.2 | 32.0 | 97.3 |
| Ligand | 134.5 | 88.5 | 112.6 | 26.6 | 10.2 | 88.7 |
| **R.m.s. deviations** |  |  |  |  |  |  |
| Bond lengths (Å) | 0.005 | 0.004 | 0.003 | 0.002 | 0.004 | 0.005 |
| Bond angles (°) | 1.299 | 1.172 | 0.620 | 0.462 | 0.602 | 1.304 |
| **Validation** |  |  |  |  |  |  |
| MolProbity score | 1.79 | 1.32 | 1.74 | 1.41 | 1.49 | 1.79 |
| Clashscore | 9.17 | 2.94 | 8.74 | 4.91 | 4.14 | 9.10 |
| Poor rotamers (%) | 0.69 | 0 | 0.37 | 0 | 0 | 0 |
| **Ramachandran plot** |  |  |  |  |  |  |
| Favored (%) | 95.70 | 96.39 | 96.06 | 97.19 | 95.77 | 95.65 |
| Allowed (%) | 4.30 | 3.61 | 3.94 | 2.81 | 4.23 | 4.35 |
| Disallowed (%) | 0 | 0 | 0 | 0 | 0 | 0 |

|  |  |  |  |  |  |
| --- | --- | --- | --- | --- | --- |
|  | R432A<br>Ca<sup>2+</sup>-closed<br>MSP2N2<br>(EMDB-41480)<br>(PDB 8TPP) | A444P<br>Ca<sup>2+</sup>-long TM6<br>(EMDB-<br>41481)<br>(PDB 8TPQ) | A444P<br>Ca<sup>2+</sup>-short<br>TM6<br>(EMDB-<br>41482)<br>(PDB<br>8TPR) | A444P<br>Ca<sup>2+</sup>-bent<br>TM6<br>(EMDB-<br>41483)<br>(PDB<br>8TPS) | A444P<br>Ca<sup>2+</sup>-long<br>TM6/short<br>TM6<br>(EMDB-<br>41484)<br>(PDB<br>8TPT) |
| **Data collection and processing** |  |  |  |  |  |
| Magnification | ×130,000 | ×105,000 | ×105,000 | ×105,000 | ×105,000 |
| Voltage (kV) | 200 | 300 | 300 | 300 | 300 |
| Electron exposure (e<sup>-</sup>/Å<sup>2</sup>) | 30.0 | 59.2 | 59.2 | 59.2 | 59.2 |
| Defocus range (μm) | -1.0 to -2.0 | -0.9 to -1.9 | -0.9 to -1.9 | -0.9 to -1.9 | -0.9 to -1.9 |
| Pixel size (Å) | 0.91 | 0.825 | 0.825 | 0.825 | 0.825 |
| Symmetry imposed | C2 | C2 | C2 | C2 | C1 |
| Micrographs | 6,368 | 11,976 | 11,976 | 11,976 | 11,976 |
| Initial particle images (no.) | 1,322,332 | 4,885,672 | 4,885,672 | 4,885,672 | 4,885,672 |
| Final particle images (no.) | 51,093 | 5,199 | 174,312 | 17,404 | 84,643 |
| Map resolution (Å) | 3.64 | 3.35 | 2.85 | 2.97 | 3.01 |
| FSC threshold | 0.143 | 0.143 | 0.143 | 0.143 | 0.143 |
| Map resolution range (Å) | 3.6-4.4 | 3.5-4.3 | 2.9-3.7 | 3.1-3.9 | 3.1-3.9 |
| **Refinement** |  |  |  |  |  |
| Initial model used (PDB code) | 6QMB | 6QMB | 6QMB | 6QMB | 6QMB |
| Model resolution (Å) | 3.68 | 3.11 | 2.90 | 2.95 | 3.06 |
| FSC threshold | 0.5 | 0.5 | 0.5 | 0.5 | 0.5 |
| Model resolution range (Å) | 3.68-4.40 | 3.11-4.30 | 2.90-3.70 | 2.95-3.90 | 3.06-3.90 |
| Map sharpening *B* factor (Å<sup>2</sup>) | -111.6 | -58.5 | -89.3 | -58.7 | -71.8 |
| Model composition |  |  |  |  |  |
| Non-hydrogen atoms | 9,118 | 9,052 | 9,456 | 9,170 | 8,908 |
| Protein residues | 1,168 | 1,154 | 1,184 | 1,176 | 1,150 |
| Ligands | 4 | 8 | 8 | 6 | 8 |
| *B* factors (Å<sup>2</sup>) |  |  |  |  |  |
| Protein | 54.0 | 86.2 | 84.7 | 108.4 | 96.1 |
| Ligand | 40.2 | 85.2 | 93.0 | 117.1 | 100.5 |
| R.m.s. deviations |  |  |  |  |  |
| Bond lengths (Å) | 0.003 | 0.003 | 0.004 | 0.004 | 0.003 |
| Bond angles (°) | 0.526 | 0.612 | 0.567 | 0.995 | 0.616 |
| Validation |  |  |  |  |  |
| MolProbity score | 1.38 | 1.81 | 1.79 | 1.81 | 1.61 |
| Clashscore | 5.34 | 9.69 | 11.57 | 10.21 | 7.81 |
| Poor rotamers (%) | 0 | 0.56 | 0.52 | 0 | 0 |
| Ramachandran plot |  |  |  |  |  |
| Favored (%) | 97.53 | 95.69 | 96.70 | 95.96 | 96.94 |
| Allowed (%) | 2.47 | 4.31 | 3.30 | 4.04 | 3.06 |
| Disallowed (%) | 0 | 0 | 0 | 0 | 0 |

## Acknowledgements

The authors thank members of the Accardi lab, Harel Weinstein and George Khelashvili for helpful discussions and suggestions. The work was supported by National Institutes of Health (NIH) grants R01GM106717 and 1R35GM152012 (to A.A.). Some of this work was performed at the Simons Electron Microscopy Center and National Resource for Automated Molecular Microscopy located at the New York Structural Biology Center, supported by grants from the Simons Foundation (SF349247), NYSTAR, and the NIH National Institute of General Medical Sciences (GM103310). Part of the work was performed at NYU Langone Health’s Cryo-Electron Microscopy Laboratory (RRID: SCR_019202) with the help of Dr. Bing Wang and Dr. William Rice, and at the Cryo-EM Core Facility at Weill Cornell Medical College with the help of Dr. Carl Fluck. Initial screening was performed at NYU Langone Health’s Cryo-Electron Microscopy Laboratory (RRID: SCR_019202) and the Cryo-EM Core Facility at Weill Cornell Medical College.

## Author contributions

Z.F. and A.A. designed the experiments; Z.F. and O.E.A. performed experiments; Z.F., O.E.A., and A.A. analyzed the data; Z.F. and A.A. wrote the paper. All authors edited the manuscript.

## Methods

### Expression and purification of nhTMEM16

Wild-type and mutant nhTMEM16 were expressed using a pYES2 vector with an N-terminal GFP-His_10_. Expression and purification of all constructs was carried out as described ^1,2^. Briefly, *S. cerevisiae* transformed with WT and mutant construct were grown in yeast synthetic drop-out medium supplemented with Uracil (CSM-URA; MP Biomedicals) to an OD_600_ of 0.8 and expression was induced with 2% (w/v) galactose at 25°C for 40 hours. Cells were collected and resuspended in lysis buffer (150 mM NaCl, 50 mM HEPES, pH 7.6) supplemented with 0.5 mM CaCl_2_, 5 μg ml^-1^ leupeptin, 2 μg ml^-1^ pepstatin and protease inhibitor cocktail tablets (Roche) and lysed with an EmulsiFlex-C3 homogenizer at pressures above 20,000 psi. Cell debris was removed by centrifugation at 3,000 g for 15 min. Membranes were subsequently harvested by centrifugation with a 45 TI rotor (Beckmann) at 200,000 g for 1 hr. Membrane proteins were extracted by supplementing the lysis buffer with 1% n-dodecyl-β-D-maltopyranoside (DDM, Anatrace), and incubated for 1.5 h at 4°C. The insoluble fraction was removed by centrifugation at 30,000 g for 30 min. The supernatant was supplemented with 10 mM imidazole, loaded onto a column of Ni-NTA agarose resin (Qiagen), washed with 150 mM NaCl, 5% (w/v) glycerol, 0.5 mM CaCl_2_, 10 mM HEPES, pH 7.6 and 0.025% DDM (Buffer A) + 50 mM imidazole and eluted with Buffer A + 400 mM imidazole. The nhTMEM16 protein peak was collected and concentrated using a 100 kDa molecular weight cut off concentrator (Amicon Ultra, Millipore). The N-terminal GFP-His_10_ tag was cleaved overnight with 3C protease before gel filtration chromatography by using a Sepharose 6 Increase 10/300 GL column (GE Healthcare). For the purification of apo proteins, 2 mM EGTA was used in all steps.

### Liposome reconstitution

Liposomes were prepared as described ^2^. A 3:1 mixture of 1-palmitoyl-2-oleoyl-snglycero-3-phosphoethanolamine (POPE) and 1-palmitoyl-2-oleoyl-sn-glycero-3-phospho-(1′-rac-glycerol) (POPG) was used for the preparation of POPE:POPG liposomes and a 7:3 mixture of 1,2-dioleoyl-sn-glycero-3-phosphocholine (DOPC) and 1,2-dioleoyl-sn-glycero-3-phospho-(1’-rac-glycerol) (DOPG) was used for the preparation of DOPC:DOPG liposomes. For the preparation of DOPE:DOPG and POPC:POPG liposomes, a 7:3 mixture of 1,2-dioleoyl-sn-glycero-3-phosphoethanolamine (DOPE) and DOPG and 1-palmitoyl-2-oleoyl-glycero-3-phosphocholine (POPG) and POPG were used, respectively. Briefly lipids in chloroform (Avanti), including 0.4% w/w 1-myristoyl-2-{6-[(7-nitro-2-1,3-benzoxadiazol-4-yl)amino]hexanoyl}-sn-glycero-3-phosphoethanolamine (NBD-PE), were dried under nitrogen gas. The dried lipids were washed with pentane and resuspended at 20 mg ml^-1^ in 150 mM KCl, 50 mM HEPES pH 7.4 with 35 mM 3-[(3-cholamidopropyl) dimethylammonio]-1-propanesulfonate (CHAPS). The proteins were added at a concentration of 5 µg protein/mg lipids. The detergent was removed by using Bio-Beads SM-2 (Bio-Rad) with rotation at 4 °C. For the mixture without POPE lipids, proteoliposomes were formed by removing the detergent with 5 times Bio-Beads exchanges of 200 mg ml^-1^. For the mixture with POPE lipids, 4 times Bio-Beads exchanges of 150 mg ml^-1^ were performed. Calcium or EGTA were introduced using sonicate, freeze-thaw cycles. The liposomes were extruded 21 times through a 400-nm membrane before use.

### Scrambling assay

The scrambling assay was carried out as described ^3^. Briefly, 20 μl of extruded liposomes were added into 2ml of external solution (300 mM KCl, 50 mM HEPES, 0.5 mM Ca(NO_3_)_2_ or 2 mM EGTA, pH 7.4). The fluorescence intensity of the NBD was monitored over time with mixing in a PTI spectrophotometer using excitation and emission wavelengths of 470 and 530 nm. After 100 s recording, sodium dithionite was added at a final concentration of 40 mM. Data was collected using the FelixGX 4.1.0 software at a sampling rate of 3 Hz.

### Quantification of scrambling activity

Quantification of the scrambling rate constants by nhTMEM16 was determined as described ^2,3^. Briefly, the fluorescence time course was fit to the following equation

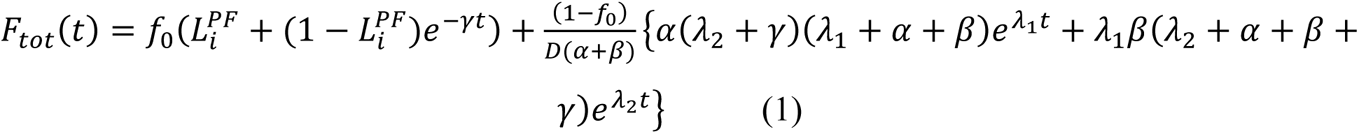

Where F_tot_(t) is the total fluorescence at time t, 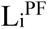 is the fraction of NBD-labeled lipids in the inner leaflet of protein free liposomes, where γ is the rate constant of dithionite reduction, f_0_ is the fraction of protein-free liposomes in the sample, α and β are respectively the forward and backward scrambling rate constants and

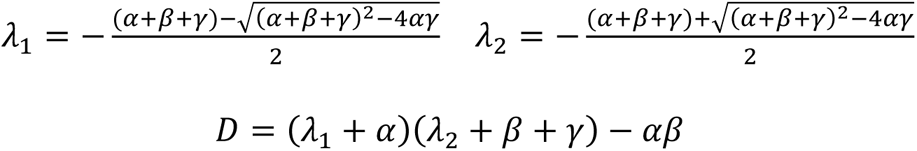

The free parameters of the fit are f_0_, α and β while 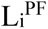 and γ are experimentally determined from experiments on protein-free liposomes. In protein-free vesicles a very slow fluorescence decay is visible, likely reflecting a slow leakage of dithionite into the vesicles or the spontaneous flipping of the NBD-labeled lipids. A linear fit was used to estimate that the rate of this process is L = (5.4±1.6)10^-5^ s^-1^ (n>160). For wild-type and mutant nhTMEM16 the leak is >2 orders of magnitude smaller than the rate constant of protein-mediated scrambling and therefore is negligible. All conditions were tested side by side with a control preparation.

### MSP1E3/MSP2N2 purification and nanodisc reconstitution

MSP1E3 and MSP2N2 were expressed and purified as described ^4^. Briefly, MSPs in a pET vector (Addgene #20064 and #29520) was transformed into BL21-Gold (DE3) strain (Stratagene). Transformed cells were grown in LB media supplemented with Kanamycin (50 mg l^-1^) until reaching an OD_600_ of 0.8. The expression of the proteins was induced by adding 1 mM IPTG (isopropyl β-D-1-thiogalactopyranoside) for 3 hours. The cells were harvested and resuspended a lysis buffer containing 40 mM Tris-HCl pH 8.0, 300 mM NaCl, 1% Triton X-100, 5 μg ml^-1^ leupeptin, 2 μg ml^-1^ pepstatin and protease inhibitor cocktail tablets (Roche). The cells were then lysed by sonication and cell debris was removed by centrifugation at 30,000 g for 45 min at 4° C. The supernatant was incubated with Ni-NTA agarose resin for one hour at 4 °C. The resin was then sequentially washed with 40 mM Tris-HCl pH 8.0 and 300 mM NaCl (Buffer B) + 1% Triton X-100; followed by a wash with Buffer B + 50 mM sodium cholate, 20 mM imidazole; and one with Buffer B + 50 mM imidazole. The protein was eluted with buffer B + 400 mM imidazole. The eluted protein was further purified by gel filtration chromatography using a Superdex 200 Increase 10/300 GL column (GE Lifesciences) pre-equilibrated with buffer C (150 mM KCl, 50 mM Tris pH 8.0) supplemented with 0.5 mM EDTA. The final protein was concentrated to ∼10 mg ml^-1^ using a 30 kDa molecular weight cut-off concentrator (Amicon Ultra, Millipore). The concentrated protein was flash-frozen and stored at −80 °C until further use.

Reconstitution of nhTMEM16 in nanodiscs was carried out as described ^5^. 7DOPC:3DOPG lipids in chloroform (Avanti) were dried under nitrogen gas, washed with pentane, and resuspended in buffer C and 40 mM sodium cholate (Anatrace) at a final lipid concentration of 20 mM. Molar ratios of 1:0.8:60 for MSP1E3:nhTMEM16:lipids and 1:0.8:150 for MSP2N2:nhTMEM16:lipids were mixed at a final lipid concentration of 7 mM and incubated at room temperature for 20 minutes. Detergent removal was carried out at 4°C via incubation with Bio-Beads SM-2 (Bio-Rad) at a concentration of 200 mg ml^-1^ for three times with two hours for each time. After reconstitution, the mixture containing the nhTMEM16-containing nanodiscs was subjected to purification using a Superose 6 Increase 10/300 GL column (GE Lifesciences) pre-equilibrated with 50 mM HEPES pH 8.0 150 mM KCl plus 0.5 mM CaCl_2_ or 5 mM EGTA and the peak corresponding to nhTMEM16-containing nanodiscs was collected for cryo-electron microscopy analysis.

### Grid preparation and data collection

3.5 uL of nhTMEM16-containing nanodiscs (3-7mg ml^-1^) supplemented with 1.5 or 3 mM Fos-Choline-8-Fluorinated (Anatrace) was applied to a glow-discharged UltrAuFoil R1.2/1.3 300-mesh gold grid (Quantifoil) and incubated for one minute under 100% humidity at 15°C. Following incubation, grids were blotted for 2 s and plunge frozen in liquid ethane using a Vitrobot Mark IV (FEI). Sample grids were screened and collected on a 300kV Titan Krios microscope (Thermo Fisher Scientific) equipped with a K3 direct detection camera (Gatan) at NYU Langone Health’s Cryo-Electron Microscopy Laboratory or 200kV Glacios microscope with a Falcon4i direct detection camera (Thermo Fisher Scientific) at the Cryo-EM Core Facility at Weill Cornell Medical College. Energy filter (either from Gatan or Thermo Fisher Scientific) set to a slit width of 20 eV was used in data collection. For additional parameters, see Table 1.

### Image processing

All datasets were processed using Relion 3.1 or Relion 4.0 ^6^. Motion correction was performed using MotionCorr2 ^6^ and contrast transfer function (CTF) estimation was carried out using CTFFIND4 ^7^ via Relion. A 2x binning was applied for motion correction in all datasets. Particle picking was conducted using a 3D volume of the nhTMEM16-nanodisc complex low-pass filtered to 20 Å. For all datasets, 2-3 rounds of 2D classification were performed to select particles that displayed structural features resembling the nhTMEM16-nanodisc complex. The selected particles underwent several iterations of 3D refinement and 3D classification, with or without alignment, to identify potential alternate conformations. Particles sorted into the same conformation were combined and subjected to 3D refinement followed by several rounds of CTF refinement ^6^ and Bayesian polishing ^8^. All 3D refinements in Relion were conducted initially without a mask, and the converged refinement was continued with a mask excluding the nanodisc. In reconstructions where no conformational heterogeneity was observed between two protomers, C2 symmetry was applied during 3D refinement to further enhance resolution. The final resolution of all maps was determined using the gold-standard Fourier shell correlation (FSC) = 0.143 criterion and applying a soft mask around the protein using Relion Postprocessing. Density modification was applied to some final reconstructions to facilitate model building by using two unfiltered half-maps with a soft mask in Phenix v1.20.1-4487 ^9,10^. Local resolution estimation for all final maps was performed using Relion. The processing strategy described here was applied to all datasets, with adaptations made for specific datasets as described below.

For wild type nhTMEM16 in the MSP1E3/+Ca^2+^ dataset, conformational heterogeneity of the groove was initially detected in 3D classifications at an early stage. 3D classes with strong and well-defined density for the closed groove were pooled and separately processed from those with poorly defined density for TM3 and TM4. The combined particles with same closed conformation were subjected to the processing strategy described above. The homogenous subset of 1,017,312 particles was used for the final 3D reconstruction, resulting in a 2.64 Å reconstruction in C2 symmetry. To achieve a better reconstruction of individual lipids, the final subset of dimeric particles was subjected to symmetry expansion and signal subtraction of individual protomers followed by 3D classification with a monomer mask on the expanded monomers. One subset containing 875,283 monomers showed strong density for lipids was selected. Local refinement followed by density modification on these particles, yielded a 2.64 Å reconstruction of the monomer. Two copies of locally refined monomers were aligned on the initial C2 dimer map and merged in Chimera using “vop maximum” command to obtain the final map. This map was used for the model building of lipids and protein. For data processing of the open state, classes with poorly defined TM3 and TM4 conformation were pooled and subjected to 5 rounds of 3D classification without alignment using a dimer mask that excluded the nanodisc density. Particles presenting the same conformation in all 5 runs were extracted using the Starparser v1.38 script (https://github.com/sami-chaaban/starparser). The resulting 55,830 particles in the open state were subjected to 3D refinement with C2 symmetry applied, resulting in a 2.85 Å reconstruction. To identify dimers with the groove in different states, symmetry expansion and single subtraction were carried out on the initially separated subset with poorly defined TM3 and TM4 and followed by 3D classification with a monomer mask on the expanded monomers. Monomers in the open or closed state were reverted to original particles and the particles present in both subsets were sorted out, resulting in 21,677 particles. 3D auto-refinement of this subset lead to a 3.40 Å reconstruction with the groove open in one protomer and closed in the other. A similar image processing strategy was also applied to the wild-type in MSP2N2/+Ca^2+^ dataset to identify the intermediate-open and open conformations.

For wild type nhTMEM16 in the MSP1E3/0 Ca^2+^ dataset, the TM6 in the final map was poorly resolved, suggesting high conformational mobility, which impeded precise model building of this region. To improve the interpretability of the map for TM6, symmetry expansion and signal subtraction followed by 3D classification were carried out with a monomer mask on a subset of 643,398 particles. This resulted in a subset of 540,140 monomers that showed strong density for TM6. 3D refinement of this subset yielded a monomer reconstruction with improved quality of TM6. A composite map was generated by merging two copies of the monomer map with the C2 dimer map using the “vop maximum” command in Chimera. This composite map further assisted in the model building of TM6.

For the A444P MSP1E3/+Ca^2+^ dataset, conformational heterogeneity of TM6 was detected in 3D classifications after the initial rounds of classification. To identify alternate conformations, the maps for individual protomers were isolated using symmetry expansion and signal subtraction. Several rounds of 3D classifications without alignment were performed using a mask only for TM1-2/5-9 on the signal subtracted protomers which yielded a subset of 388,135 particles. This resulted in a subset of monomers with TM6 in a bent conformation and Ca^2+^ density only at the lower site. The 3D reconstruction of this subset was used as a reference in subsequent masked 3D classifications, resulting in a total number of 140,913 protomers in the bent conformation. This subset of protomers was reverted to the original dimer particles, followed by the removal of duplicated particles, resulting in a subset of 17,404 particles. 3D refinement with C2 symmetry applied resulted in a 2.97 Å reconstruction. 3D classifications without alignment were also performed using a mask only for TM6-8 on the same set of signal subtracted protomers. This resulted in two subsets of protomers with distinct TM6 conformation and Ca^2+^ occupancy. One subset had a long TM6 with strong density for two Ca^2+^ sites (10.1%), and the other subset had a short TM6 with weaker density for the upper Ca^2+^ site (80.8%). These two subsets were separately selected and reverted to the original dimer particles. After removing duplicated particles, 3D refinement with C2 symmetry was carried out on the long TM6 subset, yielding a final reconstruction at 3.35 Å. For the reconstruction of the short TM6, the subset of 253,803 particles was further cleaned up by removing particles in the bent TM6 conformation. This resulted in a final subset of 174,312 particles. 3D refinement of this subset led to a 2.85 Å reconstruction with C2 symmetry applied. A similar strategy used for identifying the open/closed dimers in the wild-type in the MSP1E3/+Ca^2+^ dataset was also applied in this case, resulting a 3.01 Å reconstruction of an A444P dimer with one protomer with a long TM6 and the other with a short TM6.

Potential additional conformations were carefully examined in all datasets, especially for the wild-type in 0 Ca^2+^, R432A in MSP1E3/+Ca^2+^ and MSP2N2/+Ca^2+^ datasets that did not display heterogeneity in the initial rounds of 3D classification. We attempted classification with alignment using both global and local searches, with or without a mask excluding the nanodisc. We explored the following approaches: (*i*) varying the low-pass filter (0 to 20 Å), the T parameter (4-64), and the number of classes (4-12); *(ii)* conducting 3D classification without alignment using masks that excluded the nanodisc and varying the same parameters as the classification with alignment; *(iii)* applying symmetry expansion and signal subtraction to isolate the monomers, followed by subsequent focused classification/refinement ^11^ to account for potential alternate conformations between two protomers; *(iv)* performing classification using cryoSPARC ^12^ for all the structures described, using particles picked in Relion. In cryoSPARC, we tried 3D variability analysis on all particles selected from 2D classification, as well as on a reduced set of particles sorted using 3D classification in Relion. None of these approaches resulted in the detection of observable protein movements in these datasets.

### Model Building and Refinement

Previous nhTMEM16 structures (PDB: 6QMA, 6QMB and 6QM9) were used as starting models and fitted into the new maps using several rounds of PHENIX real space refinement ^13^. Lipids were initially modeled as POPG (PGW) and truncated according to the observed density. In most cases the full headgroup could not be resolved so the lipids were truncated at the phosphate atom. The models were inspected and areas that were better resolved were build *de novo*. The model was improved iteratively by real space refinement in PHENIX ^10^ imposing crystallographic symmetry and secondary structure restraints. Manual inspection was performed to remove the outliers. In all models, unsharpened maps were used to assist in the building process.

### Model validation

To validate the refinement, the FSC between the refined model and the final map was calculated (FSCsum). To evaluate for over-fitting, random shifts of up to 0.3 Å were introduced in the final model and the modified model was refined using PHENIX ^10^ against one of the two unfiltered half maps. The FSC between this modified-refined model and the half map used in refinement (FSCwork) was determined and compared to the FSC between the modified-refined model and the other half map (FSCfree) which was not used in validation or refinement. The similarity in these curves indicates that the model was not overfit. The quality of all three models was assessed using MolProbity ^14^ which indicates that the models are of high quality.

## Extended Data

**Extended Data Figure 1:**
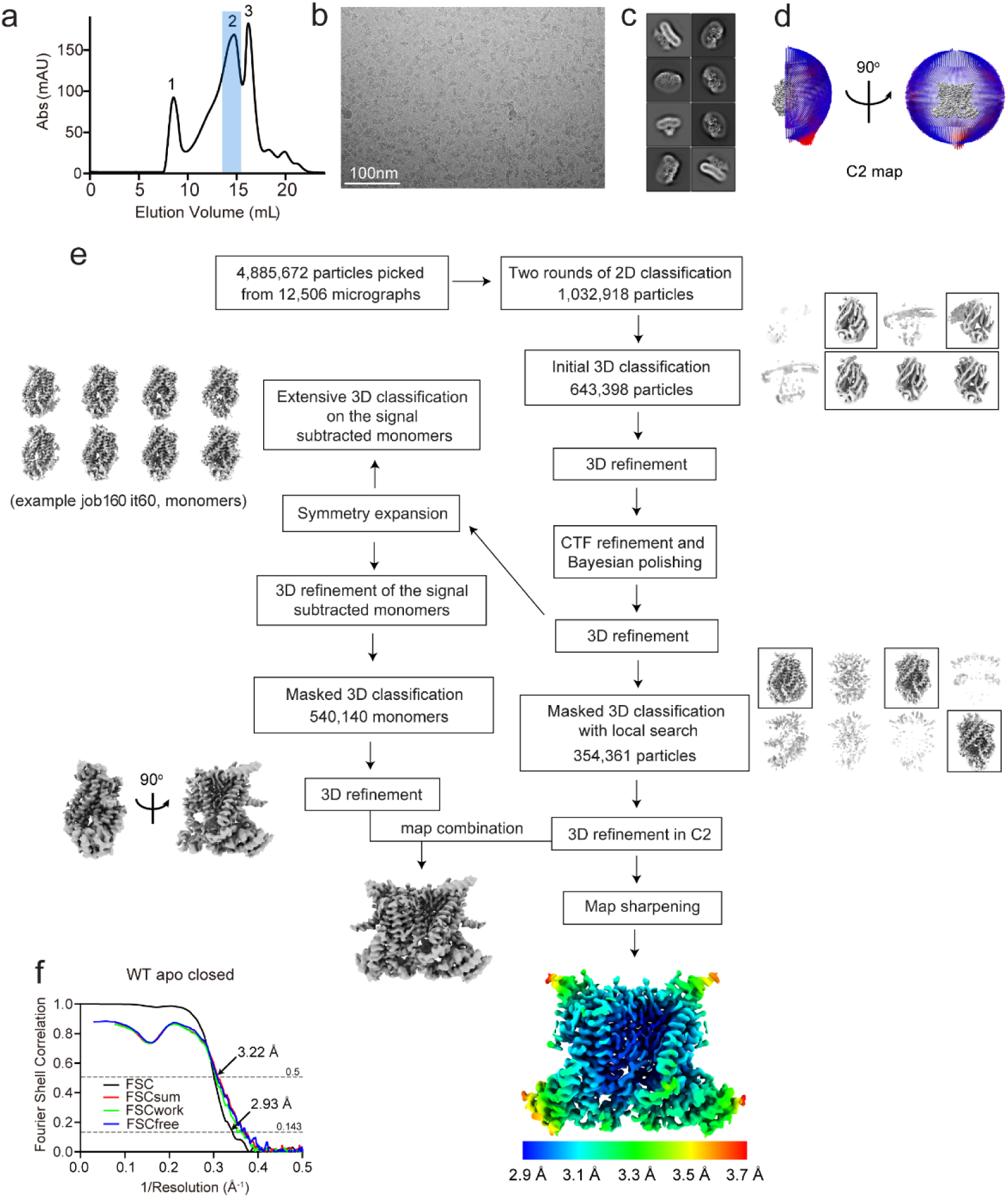
Structure determination of nhTMEM16 in the MSP1E3 nanodisc in 0 Ca^2+^. **a**, Size exclusion profile of the reconstituted nhTMEM16-nanodisc sample in 0 Ca^2+^. The peak in the blue shadow contains the nhTMEM16-nanodisc complex. **b**, Representative micrograph. **c**, Representative 2D classes of the nhTMEM16-nanodisc complex. **d**, Angular distribution of the final reconstruction with C2 symmetry. **e**, Image processing workflow including symmetry expansion and 3D classifications of the signal subtracted monomers. Final masked reconstruction colored by local resolution calculated using the Relion implementation. **f**, FSC plots for nhTMEM16-nanodisc complex in 0 Ca^2+^. FSC (black) is between the two half maps to determine the resolution of the reconstruction evaluated at 0.143 cutoff. FSCsum (red), FSCwork (green), and FSCfree (blue) are model validations evaluated at 0.5 cutoff.

**Extended Data Figure 2:**
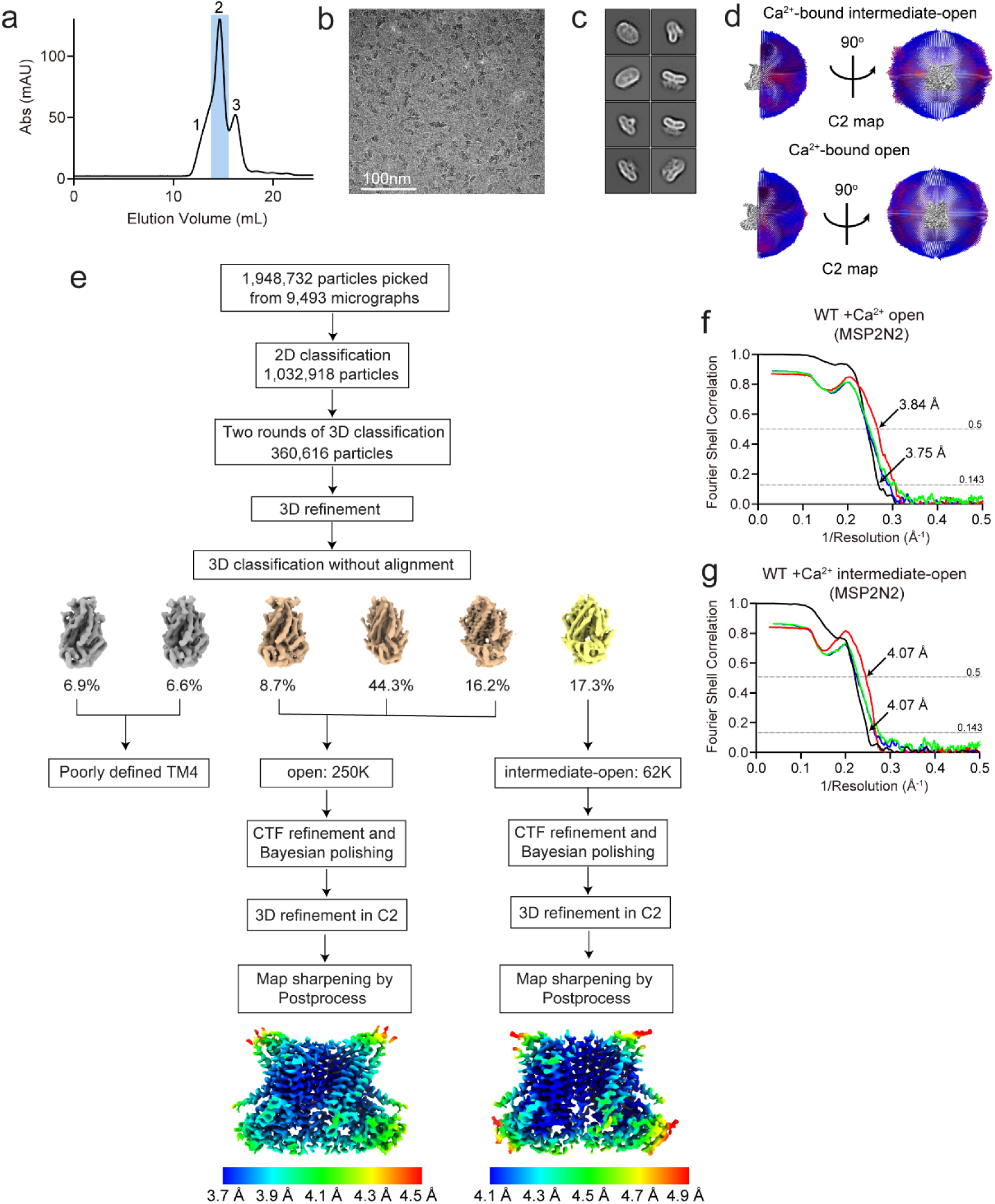
Structure determination of nhTMEM16 in the MSP2N2 nanodisc in the presence of Ca^2+^. **a**, Size exclusion profile of the reconstituted nhTMEM16-nanodisc sample in the presence of 0.5mM Ca^2+^. The peak in the blue shadow contains the nhTMEM16-nanodisc complex. **b**, Representative micrograph. **c**, Representative 2D classes of the nhTMEM16-nanodisc complex. **d**, Angular distribution of the final reconstruction of the Ca^2+^-bound intermediate-open (top) and Ca^2+^-bound open (bottom) state. **e**, Image processing workflow. Final masked reconstruction colored by local resolution calculated using the Relion implementation. **f**, **g**, FSC plots for nhTMEM16-nanodisc complex in + Ca^2+^ in the MSP2N2 nanodisc. FSC (black) is between the two half maps to determine the resolution of the reconstruction evaluated at 0.143 cutoff. FSCsum (red), FSCwork (green), and FSCfree (blue) are model validations evaluated at 0.5 cutoff.

**Extended Data Figure 3:**
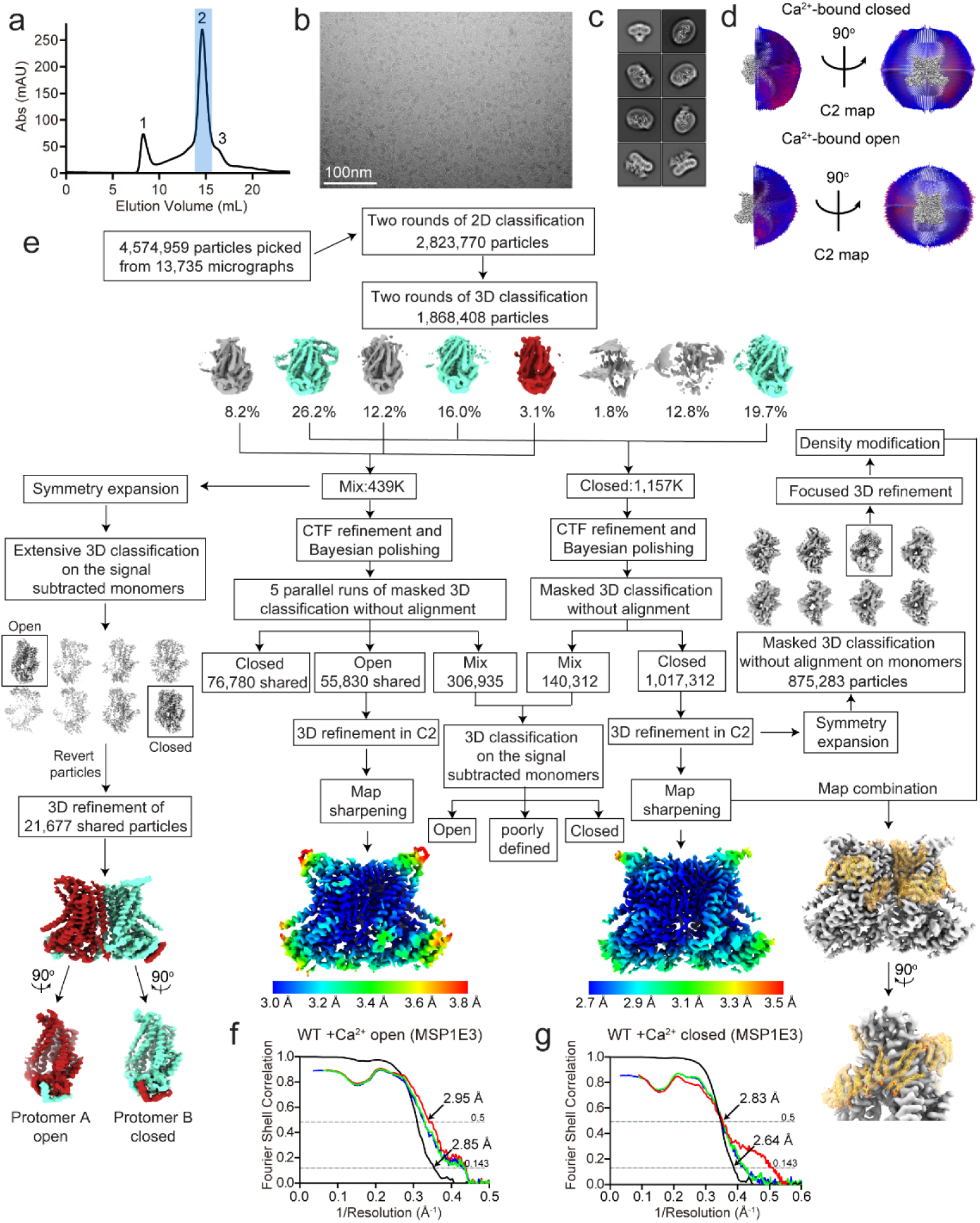
Structure determination of nhTMEM16 in the MSP1E3 nanodisc in the presence of Ca^2+^. **a**, Size exclusion profile of the reconstituted nhTMEM16-nanodisc sample in the presence of 0.5mM Ca^2+^. The peak in the blue shadow contains the nhTMEM16-nanodisc complex. **b**, Representative micrograph. **c**, Representative 2D classes of the nhTMEM16-nanodisc complex. **d**, Angular distribution of the final reconstruction of the Ca^2+^-bound closed (top) and Ca^2+^-bound open (bottom) state. **e**, Image processing workflow including symmetry expansion and 3D classifications to identify monomers with well-resolved density for lipids to assist model building, and the open/closed dimers with one open and one closed groove. Final masked reconstruction colored by local resolution calculated using the Relion implementation. **f**, **g**, FSC plots for nhTMEM16-nanodisc complex in + Ca^2+^ in the MSP1E3 nanodisc. FSC (black) is between the two half maps to determine the resolution of the reconstruction evaluated at 0.143 cutoff. FSCsum (red), FSCwork (green), and FSCfree (blue) are model validations evaluated at 0.5 cutoff.

**Extended Data Figure 4:**
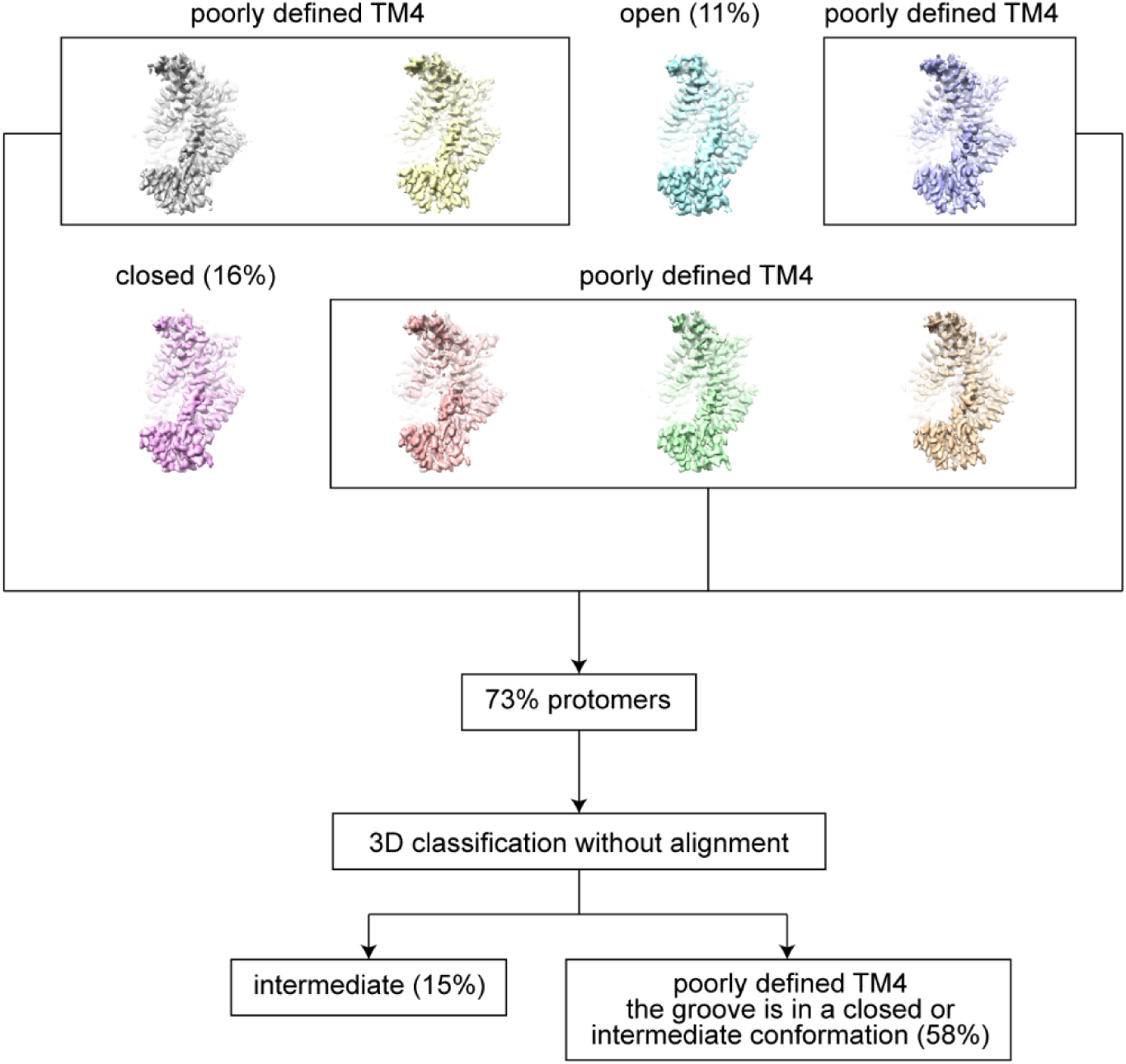
Classification of the symmetry-expanded protomers of WT nhTMEM16 in MSP1E3 nanodiscs from the particles of which the density around the groove region was not well resolved.

**Extended Data Figure 5:**
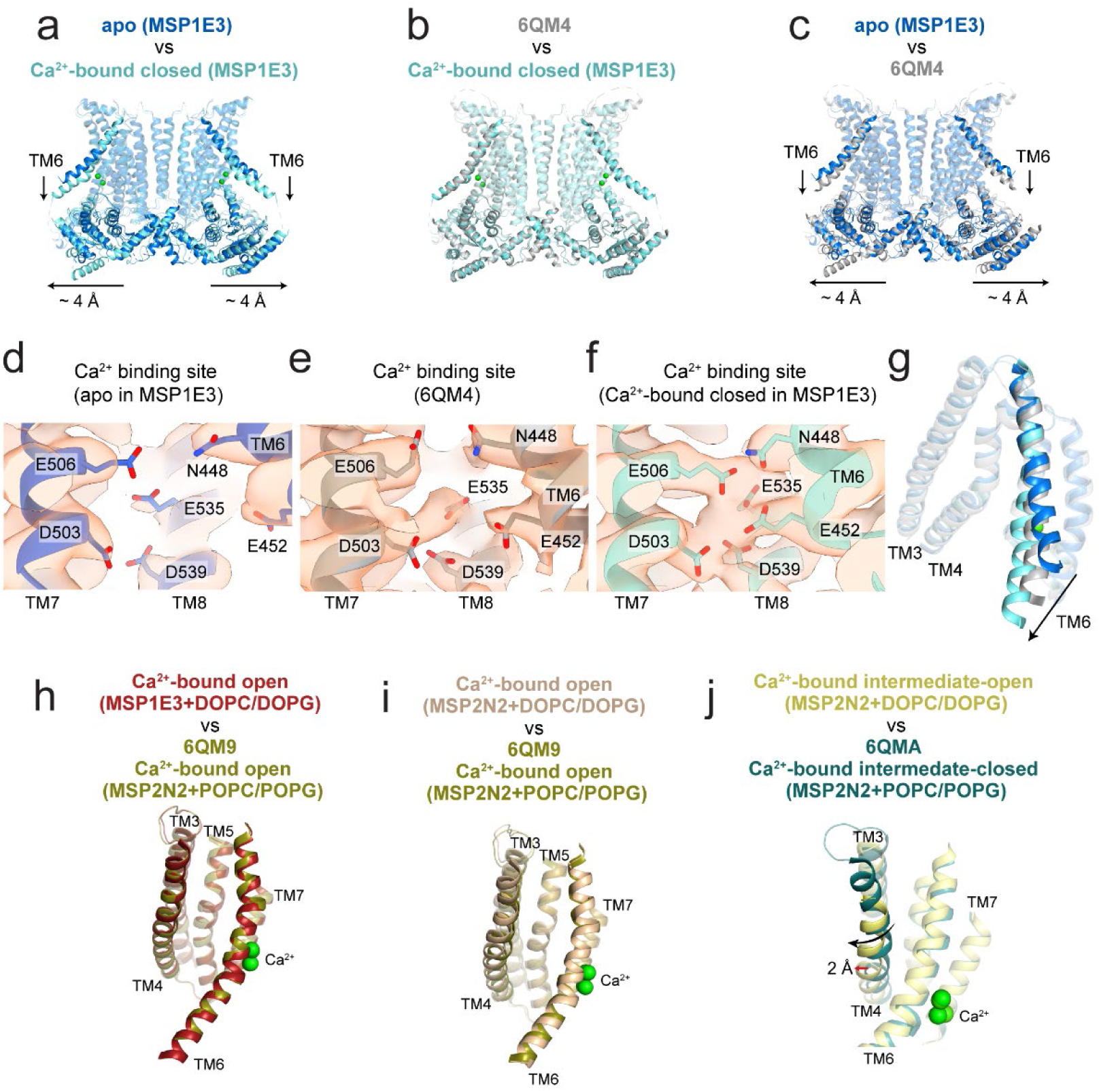
Structural characterization of the nhTMEM16 structures. **a**-**c**, Structural alignment of nhTMEM16 structures determined in DOPC/DOPG lipids and MSP1E3 nanodiscs in apo (blue) and Ca^2+^-bound closed state (cyan) to the previously reported putative apo 6QM4 conformation (gray). nhTMEM16 is viewed from the plane of the membrane from the side. Horizontal arrows at the bottom denote the ∼4 Å displacement of the cytosolic domains away from the symmetry axis between the apo state and 6QM4. Vertical arrows denote the partial straightening of TM6 from the apo state to 6QM4. Superpositions of structures in panels **a**-**c** yield a Cα r.m.s.d of 1.23 Å (**a**), 0.67 Å (**b**), and 1.76 Å (**c**), respectively. **d**-**e**, Views of the Ca^2+^-binding site in the Ca^2+^-free state of nhTMEM16 from this study (**d**) (blue), the previously reported putative apo structure (PDBID: 6QM4, gray) (**e**) and the Ca^2+^-bound closed state from this study (**f**). The density maps (contoured at 4.5 σ) are shown in orange. Weak residual density at the center of the Ca^2+^-binding site is observed in 6QM4 (**e**). **g**, Structural alignment of the nhTMEM16 groove region of the structures in apo and Ca^2+^-bound closed state from this study and 6QM4. An intermediate state of TM6 is observed in 6QM4. **h**, **i**, Comparison of the groove in the open state obtained in DOPC/DOPG lipids in MSP1E3 (**h**) or MSP2N2 (**i**) to the previously reported open structure determined in POPC/POPG lipids and MSP2N2 (PDBID: 6QM9) (olive). **j**, Comparison of the groove in the intermediate-open state in MSP2N2 (yellow) to the previously reported intermediate-closed structure (PDBID: 6QMA) (dark green).

**Extended Data Figure 6:**
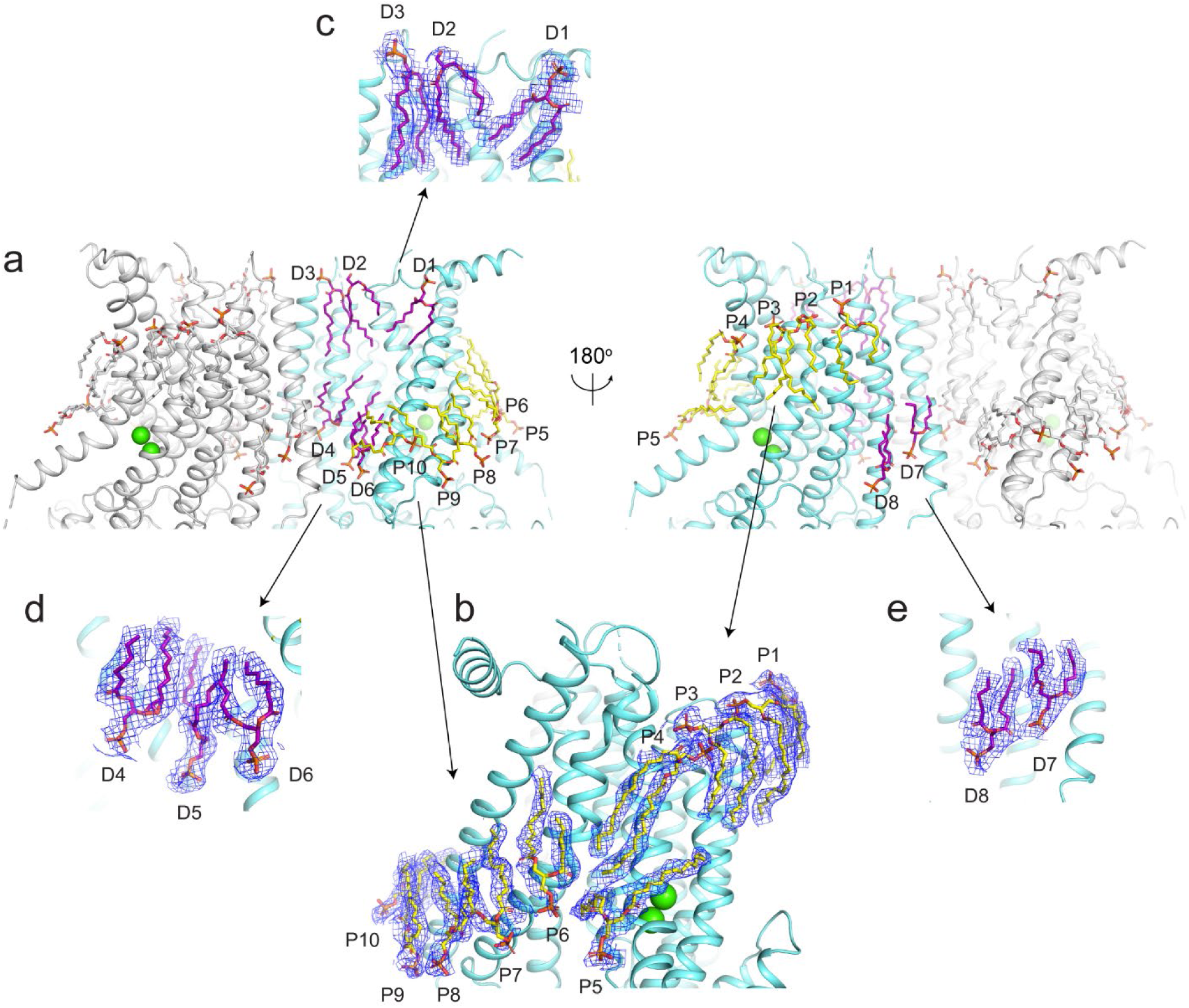
Lipid densities associated with nhTMEM16 in the Ca^2+^-bound closed state. **a**, Structure model of nhTMEM16 in the Ca^2+^-bound closed state. Protomers in the dimer are colored in gray and cyan. Acyl chains of the built lipids are colored in yellow (at the closed groove) or magenta (at the dimer cavity) in one protomer and in gray in the other one. **b**-**d**, Close-up views of lipids at the closed groove (**b**) and the dimer cavity (**c** and **d**). Lipids are shown with mesh from the sharpened map in blue with σ=2.0. Respective lipids are labelled and Ca^2+^ ions are displayed as green spheres.

**Extended Data Figure 7:**
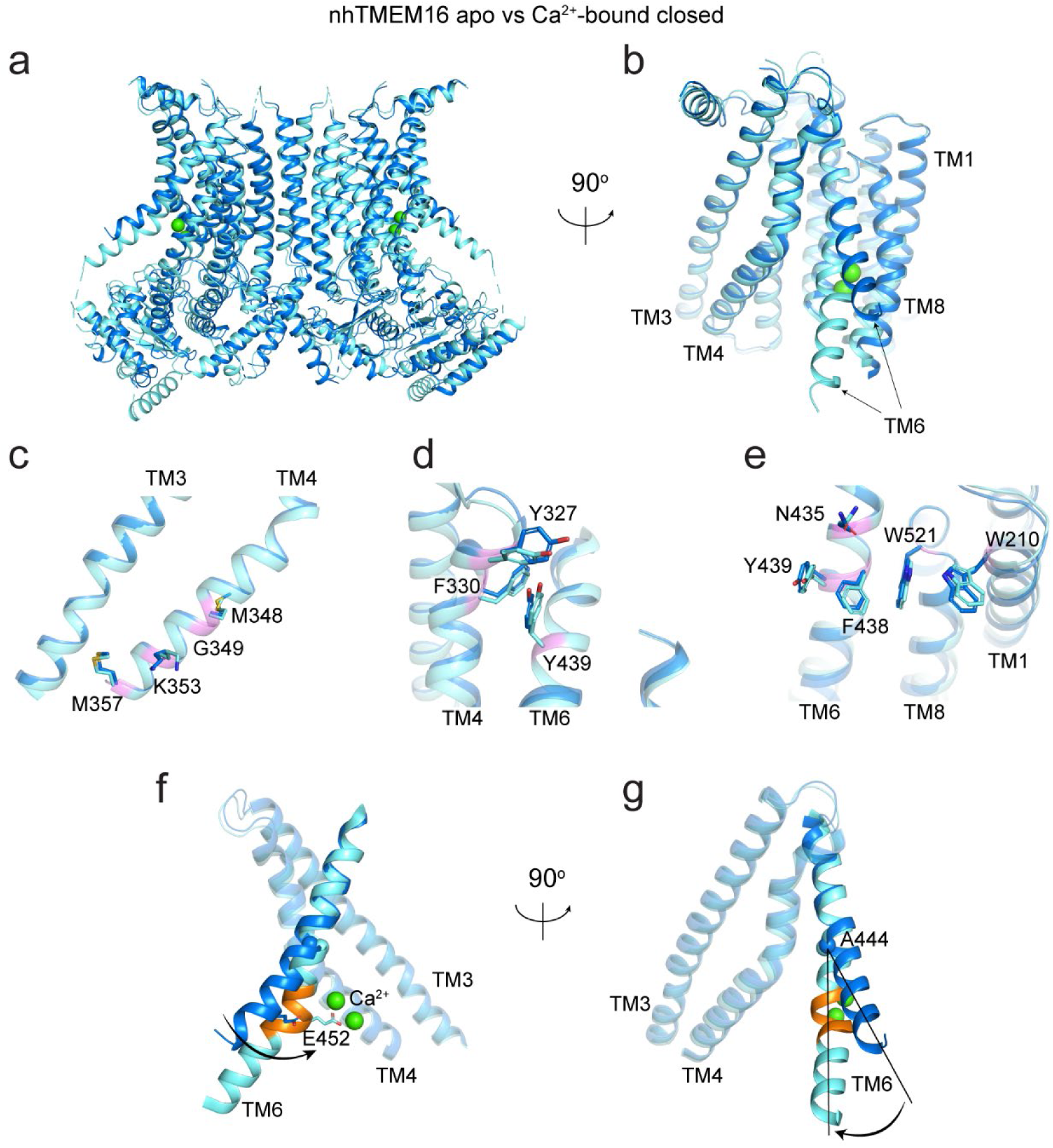
Structural comparison of nhTMEM16 apo with the Ca^2+^-bound closed state. **a**, Alignment of the apo (blue) with the Ca^2+^-bound closed nhTMEM16 (cyan). **b**, Close-up view of the groove. Ca^2+^ ions are displayed as green spheres. **d**-**e**, Close-up views of the alignment of the residues coordinating lipids outside the groove in the Ca^2+^-bound state with the equivalent residues in the apo state. Side chains are shown as sticks. Representative transmembrane helices are labelled. **f**, **g**, Conformational arrangements on the TM6 when transits from apo state (blue) to the Ca^2+^-bound closed state (cyan). π-helical turn is colored in orange.

**Extended Data Figure 8:**
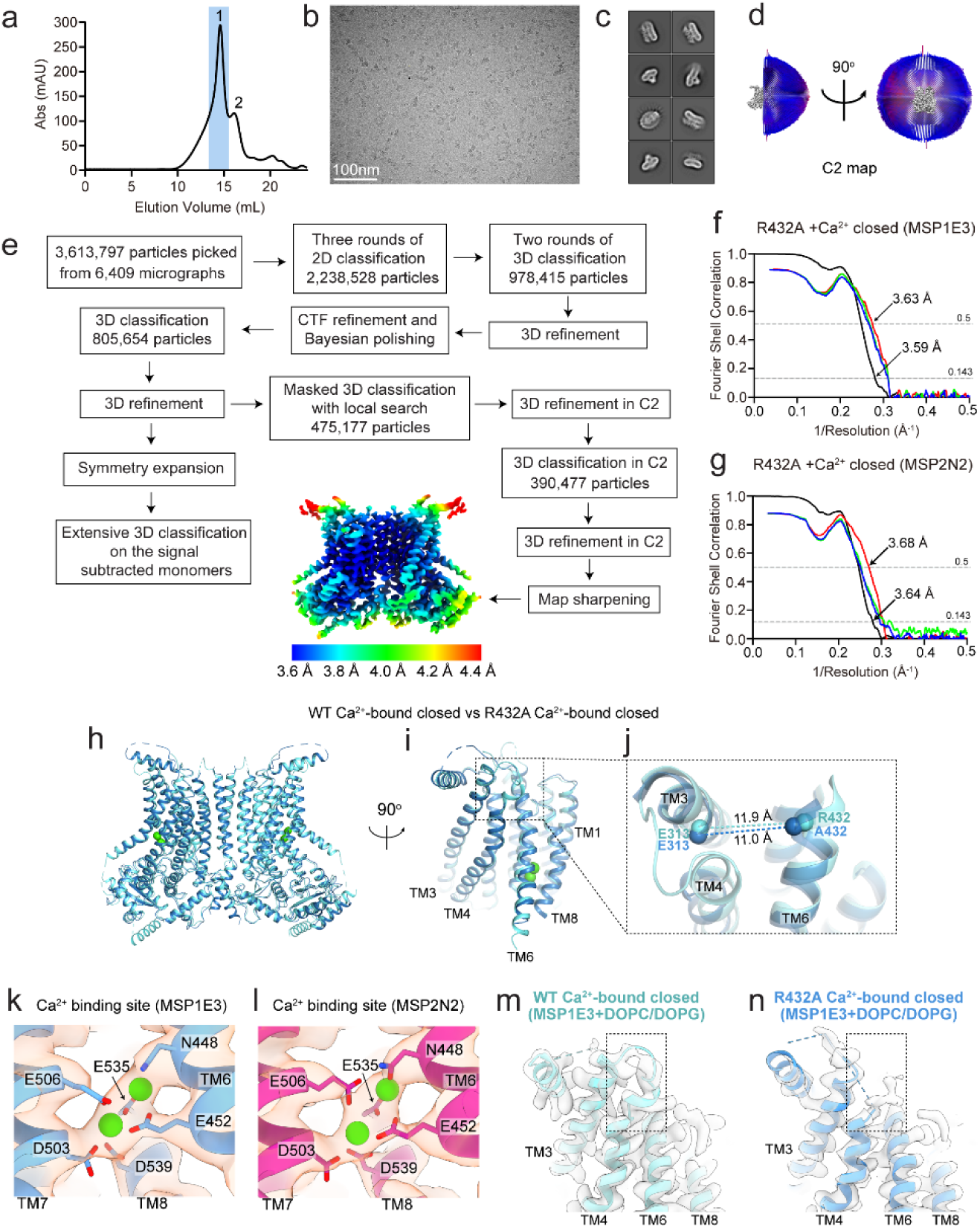
Structure determination and characterization of R432A nhTMEM16 in the MSP1E3 or MSP2N2 nanodisc in the presence of Ca^2+^. **a**, Size exclusion profile of the reconstituted the R432A nhTMEM16-nanodisc sample in the presence of 0.5mM Ca^2+^. The peak in the blue shadow contains the R432A nhTMEM16-nanodisc complex. **b**, Representative micrograph. **c**, Representative 2D classes of the R432A nhTMEM16-nanodisc complex. **d**, Angular distribution of the final reconstruction in C2. **e**, Image processing workflow including symmetry expansion and 3D classifications to identify potential alternate conformations. Final masked reconstruction colored by local resolution calculated using the Relion implementation. **f**, **g**, FSC plots for R432A nhTMEM16-nanodisc complex in + Ca^2+^ in the MSP1E3 nanodisc (**f**) or MSP2N2 nanodisc (**g**). FSC (black) is between the two half maps to determine the resolution of the reconstruction evaluated at 0.143 cutoff. FSCsum (red), FSCwork (green), and FSCfree (blue) are model validations evaluated at 0.5 cutoff. **h**-**j**, Structural comparison of R432A nhTMEM16 in the MSP1E3 nanodisc (light blue) to WT nhTMEM16 in the Ca^2+^-bound closed state (cyan) from the side (**h**) or front view (**i**). Colored spheres correspond to the position of the Cα atoms of E313 on TM3 and R432 (or A432) on TM6 and their distance is indicated (**j**). Ca^2+^ ions are shown as green spheres. **k**, **i**, Views of the Ca^2+^-binding site in the R432A nhTMEM16 in + Ca^2+^ in the MSP1E3 nanodisc (**k**) or MSP2N2 nanodisc (**i**). **m**, **n**, Views of the upper groove region of nhTMEM16 in the Ca^2+^-bound closed state (**m**) and R432A nhTMEM16 in the Ca^2+^-bound closed state (**n**).

**Extended Data Figure 9:**
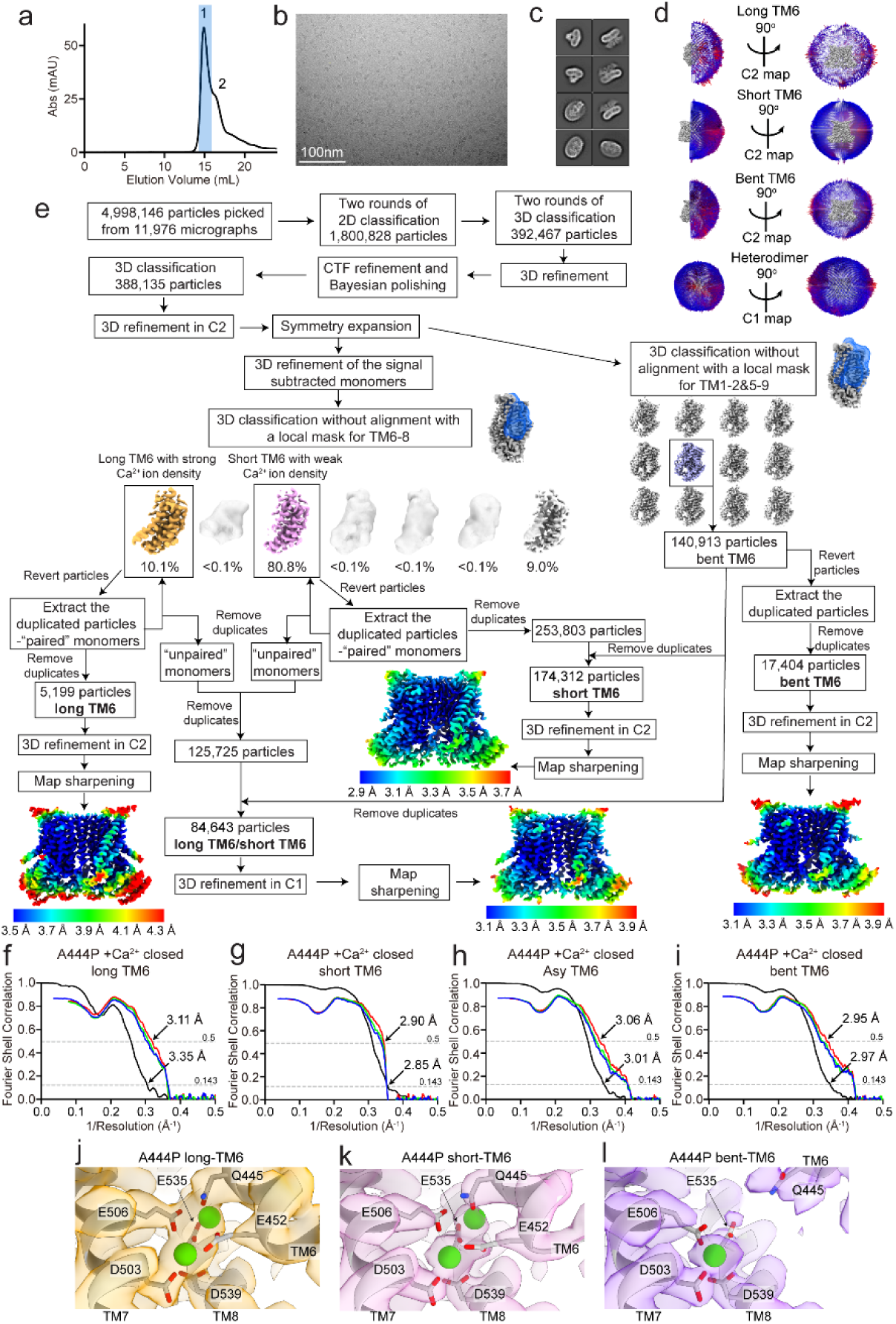
Structure determination of A444P nhTMEM16 in the MSP1E3 nanodisc in the presence of Ca^2+^. **a**, Size exclusion profile of the reconstituted the A444P nhTMEM16-nanodisc sample in the presence of 0.5mM Ca^2+^. The peak in the blue shadow contains the A444P nhTMEM16-nanodisc complex. **b**, Representative micrograph. **c**, Representative 2D classes of the A444P nhTMEM16-nanodisc complex. **d**, Angular distribution of the final reconstruction in C2. **e**, Image processing workflow including symmetry expansion and classification to identify the four different conformations, the long TM6, short TM6, bent TM6 and long TM6/short TM6 A444P. Final masked reconstruction colored by local resolution calculated using the Relion implementation. **f**-**i**, FSC plots for A444P nhTMEM16-nanodisc complex in the MSP1E3 nanodisc in the Ca^2+^-bound closed state with long TM6 (**f**), short TM6 (**g**), bent TM6 (**h**) and long TM6/ short TM6 (**i**). FSC (black) is between the two half maps to determine the resolution of the reconstruction evaluated at 0.143 cutoff. FSCsum (red), FSCwork (green), and FSCfree (blue) are model validations evaluated at 0.5 cutoff. **j**-**l**, Views of the Ca^2+^-binding site of the nhTMEM16 mutant A444P in the long TM6 state (**j**), the short TM6 state (**k**) and the bent TM6 state (**l**) with map density colored in orange, pink and purple, respectively. Side chains of the Ca^2+^ coordinating residues are shown as sticks and the Ca^2+^ ions are displayed as green spheres.

**Extended Data Figure 10:**
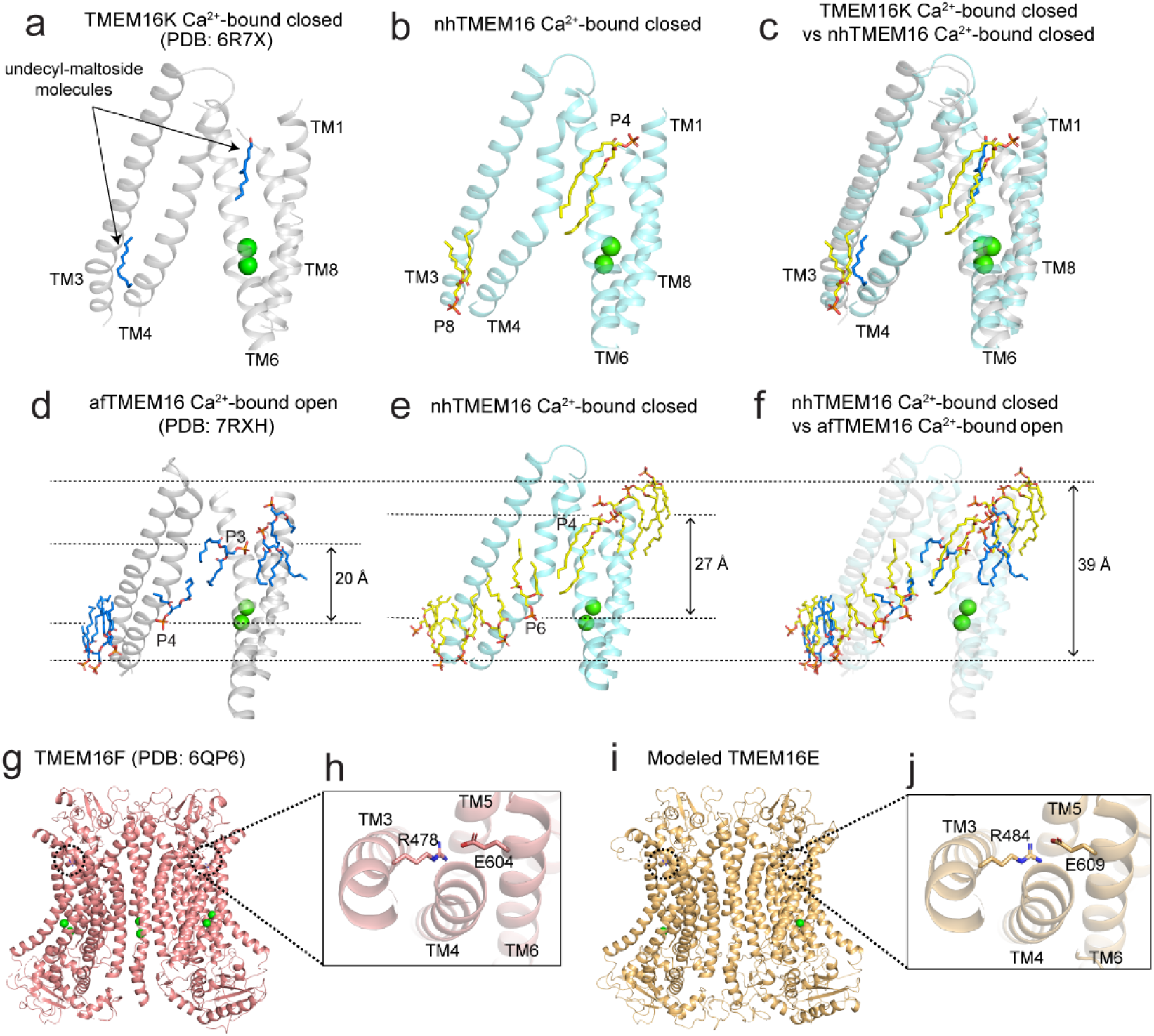
Comparison of the membrane thickness at open and closed groove and the conserved lipid/detergent binding sites and the salt bridge in mammalian TMEM16s. **a**, Two undecyl-maltoside molecules (shown in blue) were resolved in the cryo-EM structure (PDB: 6R7X, shown in gray) of TMEM16K in the Ca^2+^-bound closed state. **b**, The resolved lipids P4 and P8 associated with the closed groove of nhTMEM16. **c**, Alignment of the closed state of nhTMEM16 with TMEM16K. Ca^2+^ ions are displayed as green spheres. **d**, Lipids (shown in blue) outside of the open groove of afTMEM16 (PBD:7RXH, shown in gray). The distance between the phosphate atoms of the heads of P3 and P4 in the outer leaflet and inner leaflet is ∼20 Å. **e**, Lipids (shown in yellow) outside of the closed groove of nhTMEM16 (shown in cyan). The distance between the phosphate atoms of the heads of P4 and P6 in the outer leaflet and inner leaflet is ∼27 Å. **f**, Alignment of the rearrangements of the lipids outside of the open with the closed groove. The ∼39 Å membrane thickness is defined by the distance between the phosphate atoms of the heads of lipids away from the groove region and perpendicular to the membrane. Ca^2+^ ions are displayed as green spheres. **g, i**, The cryo-EM structure of TMEM16F (PDB:6QP6) in the Ca^2+^-bound closed state (**g**) and the predicted model of TMEM16E (**i**) by SWISS-MODEL (Waterhouse, Bertoni et al. 2018) based on TMEM16F (PDB:6QP6). The conserved salt bridge in the two structures is denoted by a dashed circle. **h**, **j**, Closed-up views of the salt bridge of R478-E604 in TMEM16F (**h**) and R484-E609 in TMEM16E (**i**).

---

**Supplementary Table 1:**
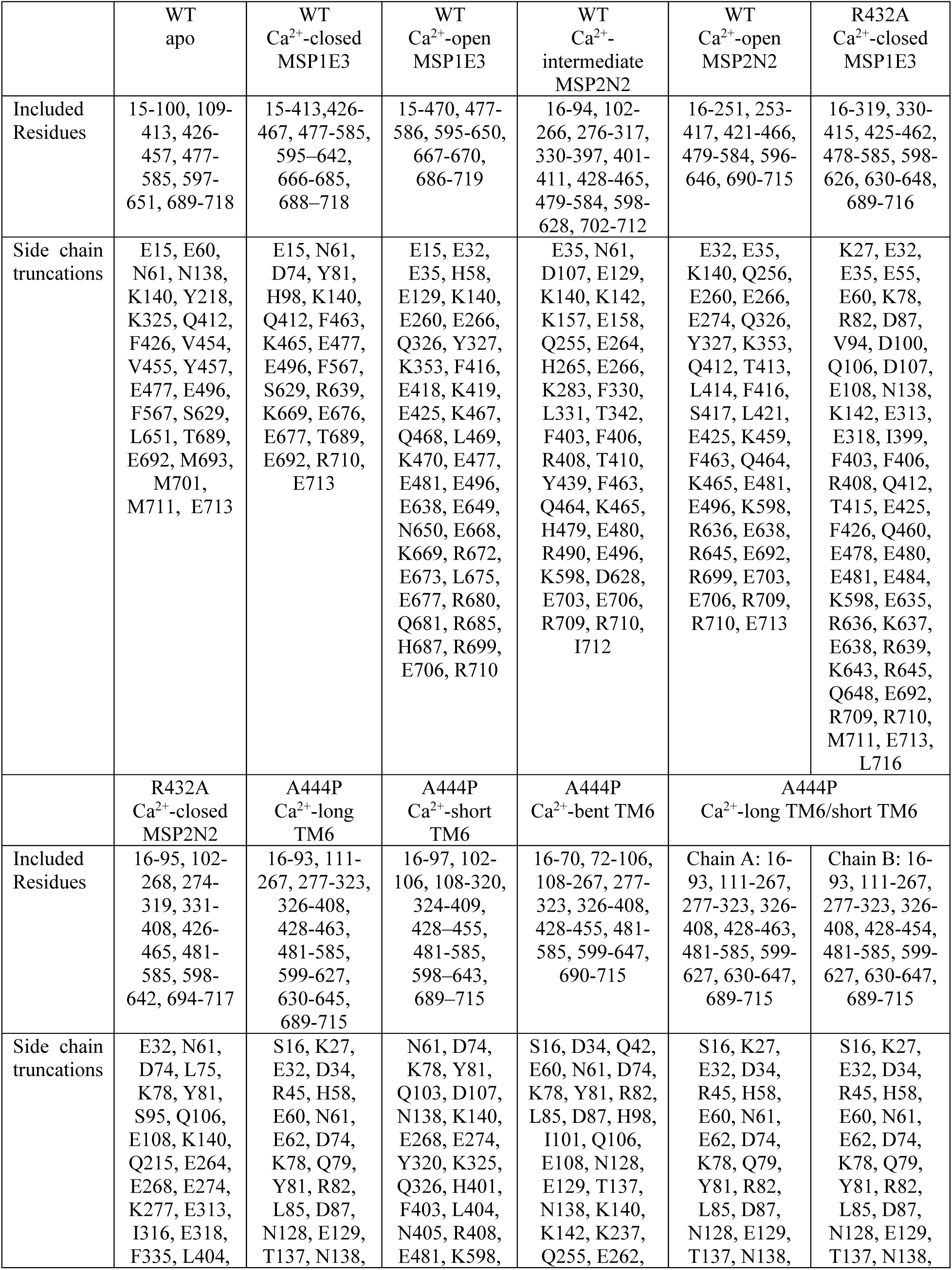
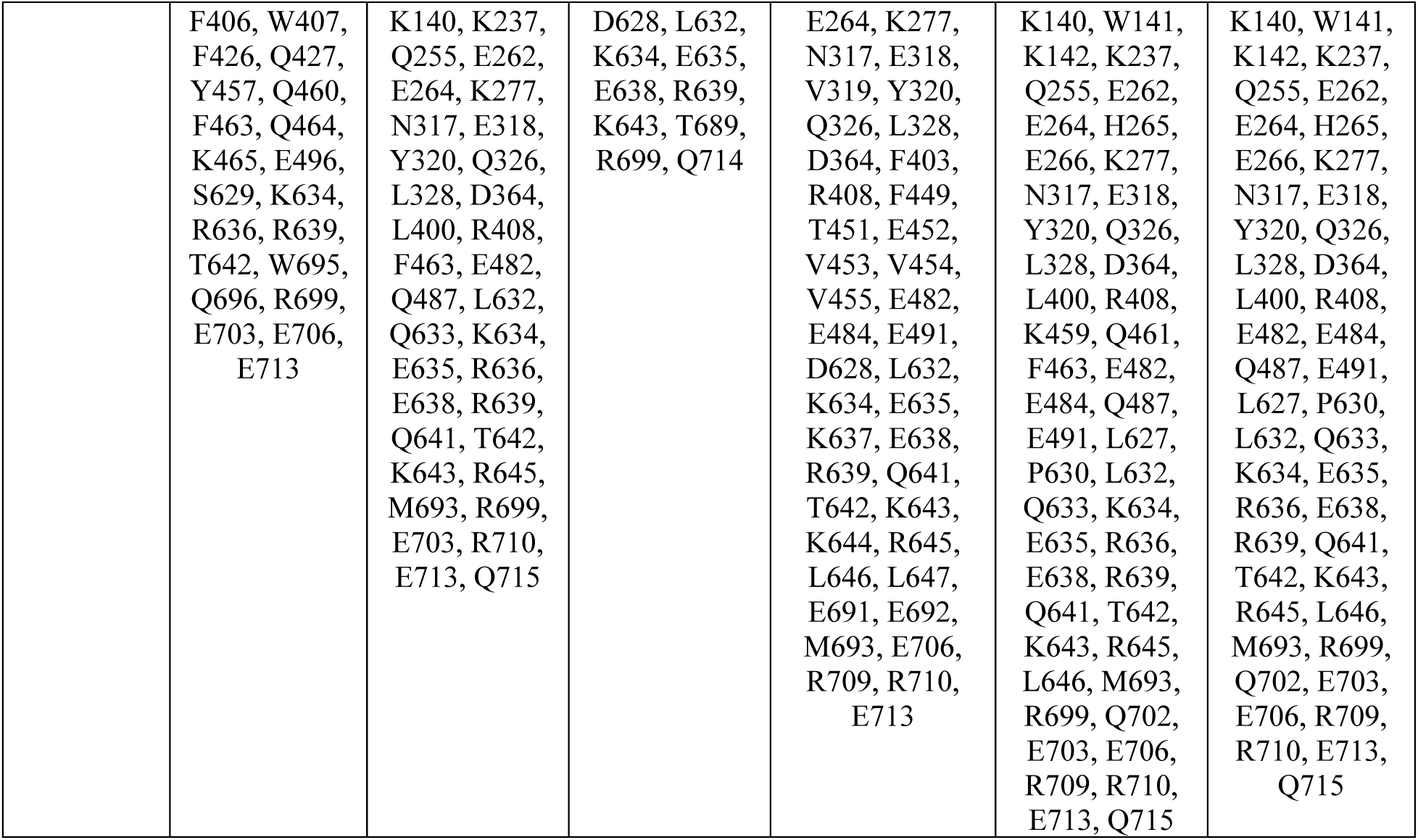
Residue ranges and side chain truncations.

