## Supplementary Table 1 for "Structural basis of closed groove scrambling by a TMEM16 protein"

**Supplementary Table 1: Residue ranges and side chain truncations**

| | WT apo | WT $Ca^{2+}$-closed MSP1E3 | WT $Ca^{2+}$-open MSP1E3 | WT $Ca^{2+}$-intermediate MSP2N2 | WT $Ca^{2+}$-open MSP2N2 | R432A $Ca^{2+}$-closed MSP1E3 |
|---|---|---|---|---|---|---|
| Included Residues | 15-100, 109-413, 426-457, 477-585, 597-651, 689-718 | 15-413,426-467, 477-585, 595–642, 666-685, 688–718 | 15-470, 477-586, 595-650, 667-670, 686-719 | 16-94, 102-266, 276-317, 330-397, 401-411, 428-465, 479-584, 598-628, 702-712 | 16-251, 253-417, 421-466, 479-584, 596-646, 690-715 | 16-319, 330-415, 425-462, 478-585, 598-626, 630-648, 689-716 |
| Side chain truncations | E15, E60, N61, N138, K140, Y218, K325, Q412, F426, V454, V455, Y457, E477, E496, F567, S629, L651, T689, E692, M693, M701, M711, E713 | E15, N61, D74, Y81, H98, K140, Q412, F463, K465, E477, E496, F567, S629, R639, K669, E676, E677, T689, E692, R710, E713 | E15, E32, E35, H58, E129, K140, E260, E266, Q326, Y327, K353, F416, E418, K419, E425, K467, Q468, L469, K470, E477, E481, E496, E638, E649, N650, E668, K669, R672, E673, L675, E677, R680, Q681, R685, H687, R699, E706, R710 | E35, N61, D107, E129, K140, K142, K157, E158, Q255, E264, H265, E266, K283, F330, L331, T342, F403, F406, R408, T410, Y439, F463, Q464, K465, H479, E480, R490, E496, K598, D628, E703, E706, R709, R710, I712 | E32, E35, K140, Q256, E260, E266, E274, Q326, Y327, K353, Q412, T413, L414, F416, S417, L421, E425, K459, F463, Q464, K465, E481, E496, K598, R636, E638, R645, E692, R699, E703, E706, R709, R710, E713 | K27, E32, E35, E55, E60, K78, R82, D87, V94, D100, Q106, D107, E108, N138, K142, E313, E318, I399, F403, F406, R408, Q412, T415, E425, F426, Q460, E478, E480, E481, E484, K598, E635, R636, K637, E638, R639, K643, R645, Q648, E692, R709, R710, M711, E713, L716 |

| | R432A $Ca^{2+}$-closed MSP2N2 | A444P $Ca^{2+}$-long TM6 | A444P $Ca^{2+}$-short TM6 | A444P $Ca^{2+}$-bent TM6 | A444P $Ca^{2+}$-long TM6/short TM6 | |
|---|---|---|---|---|---|---|
| Included Residues | 16-95, 102-268, 274-319, 331-408, 426-465, 481-585, 598-642, 694-717 | 16-93, 111-267, 277-323, 326-408, 428-463, 481-585, 599-627, 630-645, 689-715 | 16-97, 102-106, 108-320, 324-409, 428–455, 481-585, 598–643, 689–715 | 16-70, 72-106, 108-267, 277-323, 326-408, 428-455, 481-585, 599-647, 690-715 | Chain A: 16-93, 111-267, 277-323, 326-408, 428-463, 481-585, 599-627, 630-647, 689-715 | Chain B: 16-93, 111-267, 277-323, 326-408, 428-454, 481-585, 599-627, 630-647, 689-715 |
| Side chain truncations | E32, N61, D74, L75, K78, Y81, S95, Q106, E108, K140, Q215, E264, E268, E274, K277, E313, I316, E318, F335, L404, | S16, K27, E32, D34, R45, H58, E60, N61, E62, D74, K78, Q79, Y81, R82, L85, D87, N128, E129, T137, N138, | N61, D74, K78, Y81, Q103, D107, N138, K140, E268, E274, Y320, K325, Q326, H401, F403, L404, N405, R408, E481, K598, | S16, D34, Q42, E60, N61, D74, K78, Y81, R82, L85, D87, H98, I101, Q106, E108, N128, E129, T137, N138, K140, K142, K237, Q255, E262, | S16, K27, E32, D34, R45, H58, E60, N61, E62, D74, K78, Q79, Y81, R82, L85, D87, N128, E129, T137, N138, | S16, K27, E32, D34, R45, H58, E60, N61, E62, D74, K78, Q79, Y81, R82, L85, D87, N128, E129, T137, N138, |

| | | | | | | |
|---|---|---|---|---|---|---|
| | F406, W407, F426, Q427, Y457, Q460, F463, Q464, K465, E496, S629, K634, R636, R639, T642, W695, Q696, R699, E703, E706, E713 | K140, K237, Q255, E262, E264, K277, N317, E318, Y320, Q326, L328, D364, L400, R408, F463, E482, Q487, L632, Q633, K634, E635, R636, E638, R639, Q641, T642, K643, R645, M693, R699, E703, R710, E713, Q715 | D628, L632, K634, E635, E638, R639, K643, T689, R699, Q714 | E264, K277, N317, E318, V319, Y320, Q326, L328, D364, F403, R408, F449, T451, E452, V453, V454, V455, E482, E484, E491, D628, L632, K634, E635, K637, E638, R639, Q641, T642, K643, K644, R645, L646, L647, E691, E692, M693, E706, R709, R710, E713 | K140, W141, K142, K237, Q255, E262, E264, H265, E266, K277, N317, E318, Y320, Q326, L328, D364, L400, R408, K459, Q461, F463, E482, E484, Q487, E491, L627, P630, L632, Q633, K634, E635, R636, E638, R639, Q641, T642, K643, R645, L646, M693, R699, Q702, E703, E706, R709, R710, E713, Q715 | K140, W141, K142, K237, Q255, E262, E264, H265, E266, K277, N317, E318, Y320, Q326, L328, D364, L400, R408, E482, E484, Q487, E491, L627, P630, L632, Q633, K634, E635, R636, E638, R639, Q641, T642, K643, R645, L646, M693, R699, Q702, E703, E706, R709, R710, E713, Q715 |
